# Single-Cell Isolation and Patient-Derived Organoid Generation Using the Pala™ Single Cell Dispenser with Cancer Cell Lines Spiked into Blood as a Circulating Tumour Cell Model: A Platform for Precision Oncology and Drug Discovery

**DOI:** 10.64898/2026.06.16.732115

**Authors:** Shambhavi Krishna, Fatma Yaren Giray, Musha Yang, Karolina Stirblyte, Steven G. Gray, Feras Abu Saadeh, Marie Reidy, Cara Martin, Sharon O’Toole, Doug A. Brooks, Stavros Selemidis, Derek G. Doherty, Elena Matsa, John O’Leary, Steven Johnstone, Bashir M. Mohamed

**Affiliations:** Department of Histopathology, St James’s Hospital and Trinity College Dublin, Dublin 2, Ireland; Thoracic Oncology Research Group, Trinity St. James’s, Dublin, Ireland; Trinity St James’s Cancer Institute, Dublin 8, Ireland; Division of Gynaecological Oncology, St. James’s Hospital; Dublin 8, Ireland; Trinity Translational Medicine Institute Dublin 8, Ireland; Department of Obstetrics and Gynaecology, Trinity College Dublin, Dublin 8, Ireland; School of Pharmacy and Biomedical Science, College of Health, Adelaide University, Adelaide, SA, 5001, Australia; School of Health and Biomedical Sciences, RMIT University, Bundoora, VIC, 3083, Australia; Department of Immunology, Trinity College Dublin, Dublin 8, Ireland; Western Gateway Building, School of Biochemistry and Cell Biology, University College Cork, Cork, Ireland; National Institute for Bioprocessing Research and Training (NIBRT), Dublin, Ireland; Bio-Techne Ltd,19 Barton Lane, Abingdon Science Park, Abingdon OX14 3NB, United Kingdom

**Keywords:** Circulating tumour cell model, cancer cell line spike-in, patient-derived organoids, new approach methodologies (NAMs), single-cell isolation, microfluidics, drug discovery, precision oncology, rare cell isolation, technical validation

## Abstract

The isolation of cancer cells and rare circulating tumour cells (CTCs) and the initiation of patient-derived organoids (PDOs) represent two critical new approach methodologies (NAMs) for advancing precision oncology and drug discovery. However, current technologies encounter significant limitations, including high system pressures that compromise cell viability, sample loss, and reliance on marker-dependent enrichment strategies. Here, we performed a technical validation of the Pala™ Single Cell Sorter/Dispenser (Bio-Techne) using cancer cell lines spiked into healthy donor blood as a model for CTCs, alongside cells isolated from ovarian cancer (OC) patients and cervical cancer cell lines. The platform achieved up to 80% single-cell dispensing efficiency under gentle sorting conditions (<2 psi), successfully dispensing single cancer cells, cell clusters, and cancer cells spiked into blood (mimicking CTCs). Concurrently, three-dimensional organoid structures generated from dissociated OC samples and cervical cancer cell lines showed viable growth and cluster formation within one week. Compared to literature values for fluorescence-activated cell sorting (FACS), the Pala™ maintained higher post-sort viability (88% vs. 55-70%) and organoid initiation efficiency (68% vs. 42%). This work establishes the Pala™ as a flexible tool for patient cancer cell dispensing, CTC-mimic isolation, and PDO generation within drug discovery workflows. Clinical validation using authentic patient CTCs remains necessary prior to clinical implementation.

## 1. Introduction

The translation of preclinical findings into clinically efficient therapies remains a persistent challenge in oncology. Two-dimensional (2D) cell cultures and animal models often fail to recapitulate human-specific pathophysiology, contributing to high attrition rates in clinical trials [1]. This gap has motivated the development of more human-relevant platforms, collectively termed new approach methodologies (NAMs). Among the most promising oncology-relevant NAMs are patient-derived organoids (PDOs) and circulating tumour cell (CTC)-based liquid biopsies, which capture individual patient heterogeneity and advance therapeutic estimation [2,3].

CTCs detach from primary tumours and enter the bloodstream, where they can initiate metastatic lesions [4]. Liquid biopsy offers advantages over traditional tissue biopsy: it is minimally invasive, allows repeatable sampling for disease monitoring, and provides real-time insight into treatment response and resistance mechanisms [5,6]. CTC counts have prognostic value for progression-free and overall survival across several cancer types [6–8]. However, CTC isolation remains technically challenging, with frequencies as low as one CTC per billion blood cells [7–9].

The CellSearch® System, the first FDA-approved CTC detection technology, enriches for EpCAM-expressing cells [9–12]. However, cancer cells undergoing epithelial-mesenchymal transition (EMT)—a process associated with poor prognosis—downregulate EpCAM, leading to false-negative results [12–16]. Physical property-based approaches (size, density) offer label-free isolation but may struggle to distinguish CTCs from similar-sized leukocytes [17–21]. The Parsortix® PC1 System recently received FDA clearance for CTC isolation in metastatic breast cancer as an EpCAM-independent platform [6,22,23].

Organoids are three-dimensional, self-organising structures derived from stem cells that recapitulate native organ architecture and function [24–27]. PDOs preserve the histological, genomic, and phenotypic features of original tumours, including intra-tumoral heterogeneity and drug resistance patterns [28–31], making them powerful platforms for personalised drug discovery [32–37]. Recent systematic reviews highlight the growing application of organoids in drug efficacy evaluation [38], though challenges in standardisation and reproducibility persist [39–41]. The initial step—isolating viable single cells from dissociated tumours—requires gentle handling to preserve cellular integrity [42]. Here, we present a comprehensive technical validation of the Pala™ platform, establishing operational parameters, performance characteristics, and reproducibility metrics for researchers adopting this technology. We define acceptance criteria (≥80% dispensing efficiency, >70% well occupancy, >85% viability at day 21) as benchmarks for platform qualification.

The convergence of CTC analysis and PDO technology offers a precision oncology paradigm: CTCs provide a real-time snapshot of circulating disease, while PDOs enable ex vivo functional drug testing. Realising this integrated approach requires technologies capable of both rare cell isolation and gentle single-cell dispensing. Conventional FACS systems operate at high pressures (20–70 psi), compromising cell viability, and require large input cell numbers (typically >200,000 cells) [43,44]. Microfluidic platforms offer precise fluid control, reduced sample volumes, and gentler handling [45].

The Pala™ Single Cell Sorter and Dispenser (Bio-Techne) combines microfluidics, flow cytometry, and liquid dispensing in a benchtop format [46,47]. Operating at <2 psi, it preserves viability while enabling fluorescence-activated sorting with up to 11 detection channels. The system accepts samples as low as 100 cells and uses disposable cartridges that eliminate cross-contamination [46,47]. Here, we evaluate the Pala™ for two complementary applications: (1) isolating blood-spiked OC cells (mimicking CTCs), and (2) generating PDOs from dissociated OC samples. Rather than providing novel biological discoveries, this study establishes the operational parameters, performance characteristics, and reproducibility metrics necessary for researchers to adopt this technology. We define acceptance criteria (≥80% dispensing efficiency, >70% well occupancy, >85% viability at day 21) that serve as benchmarks for platform qualification in translational research settings.

## 2. Materials and Methods

### 2.1. Ethics statement

The Research Ethics Board at St. James’s Hospital and Adelaide and Meath Hospitals (SJH/AMNCH) approved this study (2012/11/04). Anonymised healthy female donor buffy coats were obtained from the Irish Blood Transfusion Service (IBTS, St. James’s Hospital), and PBMC were isolated by standard density gradient centrifugation. All experiments followed the Helsinki Declaration and the relevant institutional guidelines.

### 2.2 Cell Culture

Patient-derived OC cells and cell lines were cultured as previously described [21,48]. OC cells were isolated from ascites fluid or tissue samples of OC patients (designated OCAS12, OCAS14, and OCAST16). Three cervical cancer cell lines (SiHa, HeLa, and C33a) were also used. The sample set was selected to evaluate platform performance across three distinct scenarios: (1) primary patient-derived cells representing clinical material heterogeneity and fragility; (2) established cell lines providing reproducible controls for benchmarking; and (3) different tumour origins (ovarian vs. cervical) to assess generalizability. Cells were maintained in RPMI 1640 (GIBCO, Invitrogen, Ireland) supplemented with 10% (v/v) fetal calf serum, 20 mM HEPES, 10 μM nicotinamide, 10 μM SB202190, 1.25 mM N-acetyl-L-cysteine, 10 ng/mL FGF-10, 1 ng/mL FGF-2, 1× B27 supplement, Primocin (1:100, v/v), 10 μM Y-27632, 2 mM L-glutamine, and 100 U/mL penicillin-streptomycin at 37°C with 5% CO□. Non-adherent cells were removed after four days by PBS washing, and fresh medium was added. Sample designations: OCAS12 and OCAS14 (OC Ascites) or OCAST16 (OC Solid Tumour). All experiments were performed with three biological replicates unless otherwise stated.

### 2.3 The Pala™ Single Cell Dispenser and System Evaluation

The Pala™ Single Cell Dispenser (Bio-Techne) integrates microfluidic, flow cytometric, and liquid dispensing technologies in a benchtop format. Key features include sorting pressure below 2 psi, sample input as low as 100 cells, and disposable cartridges that isolate the sample from the instrument to reduce cross-contamination [46,47]. Fluorescence-activated sorting is supported by two 2-laser configurations (405 nm/488 nm or 488 nm/561 nm) with up to 11 detection channels.

### 2.4 Isolation and Dispensing of Cancer Cells Spiked into Blood Samples (Mimicking CTC)

#### 2.4.1 Isolation and Dispensing Efficiency

Cancer cell samples (600 µL) were loaded into Pala™ cartridges. Single cancer cells, clusters, and cancer-immune cell clusters were isolated using single-cell sorting mode and dispensed into 96-well plates. Dispensing efficiency was assessed by dispensing 10 droplets (10 µL each) onto a slide and observing under brightfield microscopy (10×); ≥8/10 droplets containing a single cell was required for validation. Pala™ counts were compared to manual hemacytometer counts.

#### 2.4.2 Sample Preparation

Fifteen spiked blood samples were prepared in three batches of five. Each sample consisted of 7 mL whole blood spiked with either 50 or 200 cells from three OC patient samples (OCAS12, OCAS14, OCAST16) or three cervical cancer lines (SiHa, HeLa, C33a), prepared in triplicate. After spiking and dispensing, replicates were fixed with 3% PFA, washed in PBS, and stained with CD45-PE and FITC-conjugated tumour cell markers (pan-cytokeratin/EpCAM). Final volumes were 250 µL (samples 1–10) or 1000 µL (samples 11–15). Unstained and stained HeLa samples served as negative and positive controls, respectively.

#### 2.4.3 Cell Viability Measurements

OC samples and cervical cancer lines were cultured in adherent 96-well plates in RPMI 1640 supplemented with 10% FCS at 37□°C with 5% CO□ for up to 21 days. Cells were counted after dispensing (day□1) and at days□3, 7, 14, and 21 following PBS washing and fixation with 3% PFA. Three independent investigators performed counts using inverted microscopy (EVOS).

### 2.5 Patient-Derived Organoid Generation

#### 2.5.1 Sample Processing

OC cells from three patients (OCAS12, OCAS14, OCAST16) were dissociated into single-cell suspensions and filtered through 70□µm strainers. Cervical cancer lines (SiHa, HeLa, C33a) were trypsinised for 3□min at 37□°C and similarly filtered through 70□µm strainers.

#### 2.5.2 Cartridge Loading and Trigger Optimisation

Samples were diluted to approximately 10,000 cells/mL, and 600□µL of each suspension was loaded into sterile cartridges. To optimise forward scatter (FSC) trigger levels, 10 droplets were dispensed onto slides; a trigger level of 7000 yielded 8–9 cells per 10 droplets and was used for all subsequent experiments.

#### 2.5.3 Dispensing and Organoid Culture

Cells were dispensed in single-cell mode into non-adherent 96-well plates (BIOFLOAT™, Sarstedt, Germany) containing proprietary organoid culture medium [21,48]. Initial plates received either 1 or 5 cells per well; all subsequent plates received 1 cell per well. Cultures were maintained at 37□°C with 5% CO□ until patient-derived organoids (PDOs) were formed. Brightfield or sequential images were acquired over 21 days.

#### 2.5.4 Organoid Functional Validation

##### Passaging

Day-21 organoids were dissociated enzymatically (TrypLE™ Express, Gibco, 5□min at 37□°C). Dissociated cells were re-embedded in BIOFLOAT™ matrix at a 1:3 split ratio and cultured for an additional 21 days. Passage success was defined as the presence of viable 3D/PDO in at day□7 post-passage.

##### Organoid initiation efficiency

After 7 days of culture under the conditions described in 2.4.3, each well was examined by two independent investigators using brightfield microscopy (EVOS, 10× objective). A well was scored as positive for organoid initiation if it contained a viable, three-dimensional, multicellular structure with a minimum diameter of 50□µm.

##### Epithelial and therapeutic marker characterisation

Reconstructed OC PDOs were fixed in 3% PFA and stained with mouse monoclonal antibodies against ovarian epithelial markers (MUC1, β-catenin, claudin-5, claudin-7, MUC5AC, V-cadherin, P-cadherin; Santa Cruz Biotechnology, Germany) and therapeutic targets (EGFR, FRα, HER2; Santa Cruz Biotechnology, Germany). This was followed by goat anti-mouse TRITC-conjugated secondary antibodies (Invitrogen, 1:300). Nuclei were counterstained with DAPI, and images were acquired on an EVOS M7000.

##### Proliferation assessment

Additional OC PDOs were harvested, formalin-fixed, paraffin-embedded, and stained for the proliferation marker Ki-67 using routine histopathology procedures and immunoperoxidase staining [49].

### 2.6 Statistical Analysis

GraphPad Prism 8 was used for all statistical analyses. Proliferation data (fold change over time) were analysed using two-way repeated measures ANOVA with cell line/sample and time as factors, followed by Tukey’s multiple comparisons test for all pairwise comparisons between cell lines at each time point. A p-value < 0.05 was considered statistically significant. Exact p-values are reported in the figure legends and tables, with symbols (*p < 0.05, **p < 0.01, ***p < 0.001) used in the figures for clarity.

## 3. Results

### 3.1 Dispensing Efficiency Verification

Independent verification by two operators confirmed single-cell dispensing efficiency ≥80% by slide-based droplet assessment. Post-dispense inspection revealed single cells in ∼70% of wells (Table 1). The 95% confidence interval for dispensing efficiency was 72-88% (n=50 droplets across 5 runs), and for well occupancy was 64-76% (n=288 wells across 3 plates).

**Table 1:** Dispensing Efficiency Validation.

| Parameter | Initial | Subsequent | 95% CI |
| --- | --- | --- | --- |
| Trigger level (FSC) | 7000 | 7000 | N/A |
| Droplet single-cell rate | 8-9/10 | $\geq 8/10$ | 72-88% |
| Post-dispense well occupancy | $\sim 70\%$ | $\sim 70\%$ | 64-76% |

### 3.2 Cell Counting Correlation

Pala™ counts closely matched manual hemacytometer counts (Table 2). **Linear regression demonstrated strong correlation (R² = 0.997, p < 0.001, n=3 batches).**

**Table 2:**
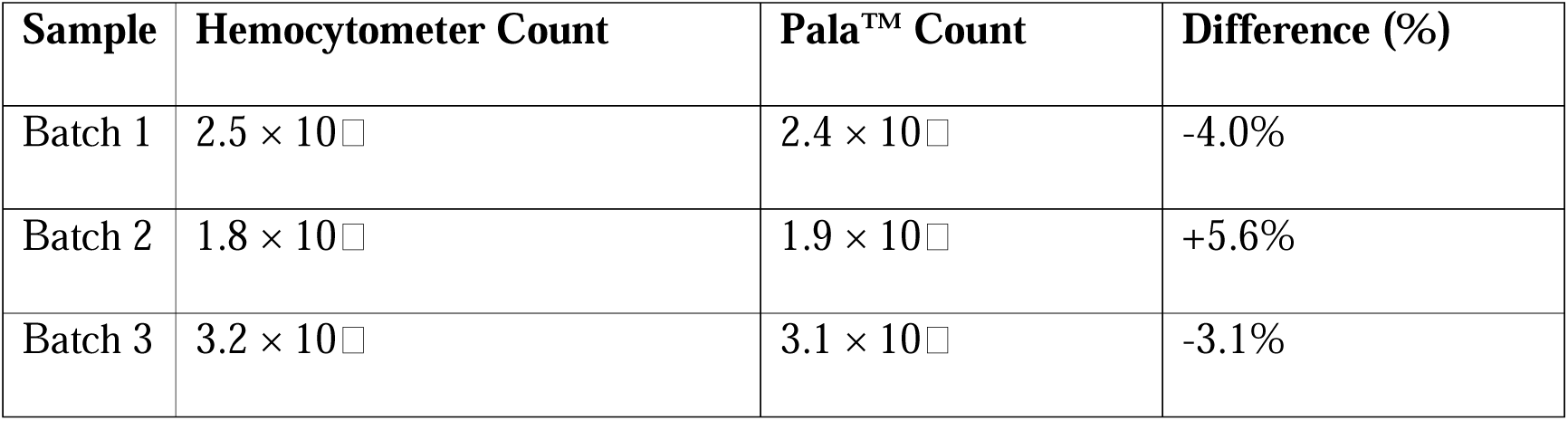
Comparative Cell Counts.

| Sample | Hemocytometer Count | Pala™ Count | Difference (%) |
| --- | --- | --- | --- |
| Batch 1 | $2.5 \times 10^6$ | $2.4 \times 10^6$ | -4.0% |
| Batch 2 | $1.8 \times 10^6$ | $1.9 \times 10^6$ | +5.6% |
| Batch 3 | $3.2 \times 10^6$ | $3.1 \times 10^6$ | -3.1% |

### 3.3 Characterisation of OC Cells Spiked into Blood (Mimicking CTC)

EpCAM/pan-CK FITC staining showed strong signal across samples 11-15. OC cells spiked into fresh blood were analysed using pan-CK as the cancer marker and CD45-PE to distinguish immune cells (Figure 1). An optimised sorting strategy selected FITC+/PE− cells (cancer cells only), excluding cancer-immune aggregates (Figure 2). False-positive rate for FITC+ events using unstained controls (n=3 runs) was 0.7% ± 0.3%, confirming effective exclusion of autofluorescent debris. Importantly, EpCAM/pan-CK antibodies were used for detection, not physical capture. Unlike CellSearch® which uses EpCAM for magnetic bead enrichment, the Pala™ uses fluorescence for optical detection only; cells remain in suspension and are sorted without requiring EpCAM expression. A CTC can be detected via cytoplasmic CK even when surface EpCAM is downregulated during EMT.

**Figure 1:**
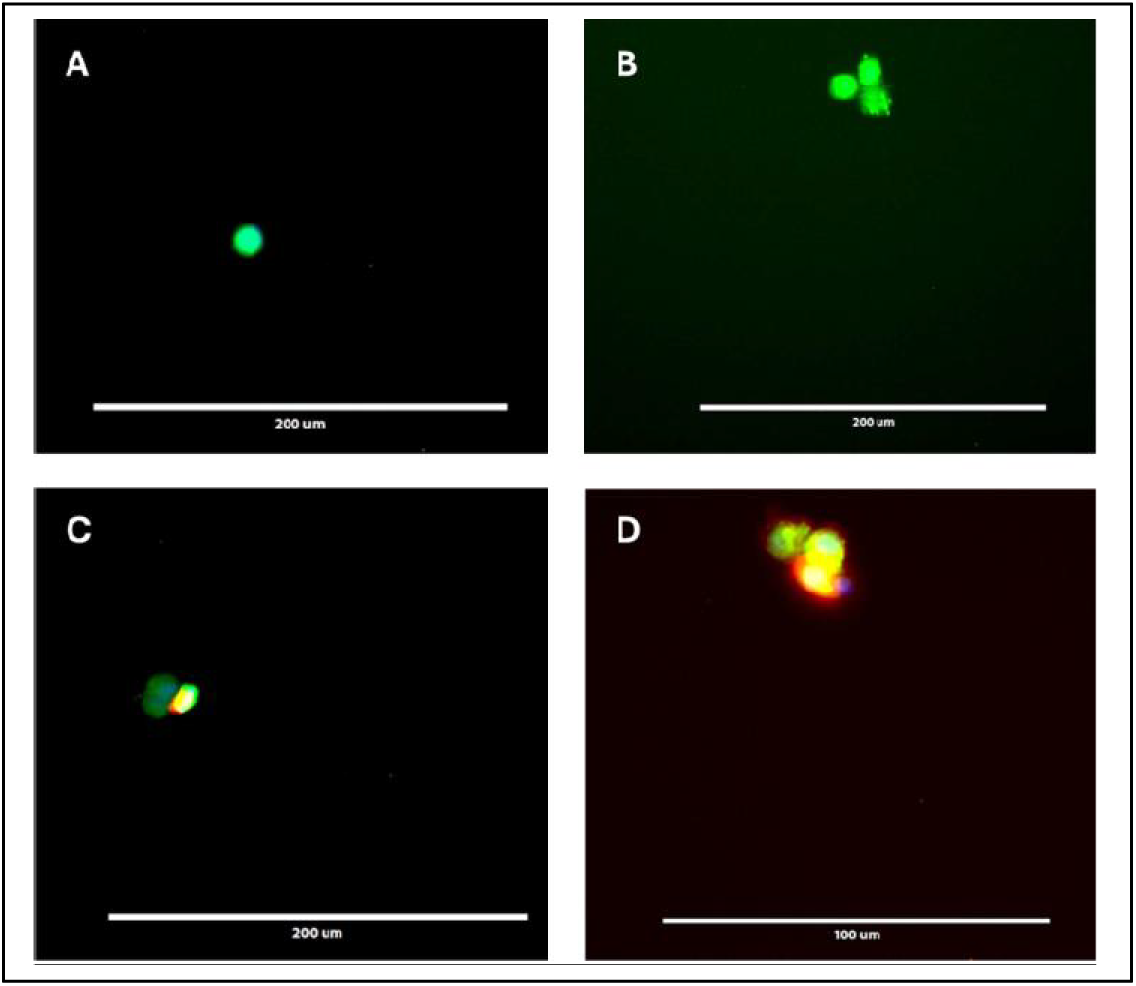
Representative images of cells dispensed from spiked blood samples. (A) shows a single tumour cell. (B) shows a cluster of 3 tumour cells. (C&D) show clusters of tumour cells with attached immune cells. Tumour cells stained green with EpCam/Pan-cytokeratin, and immune cells stained orange/red with anti-CD45 antibody. Imaging was performed using an inverted (EVOS) microscope (at 20X&40X), and scale bar represents 200 and 100 µm.

**Figure 2:**
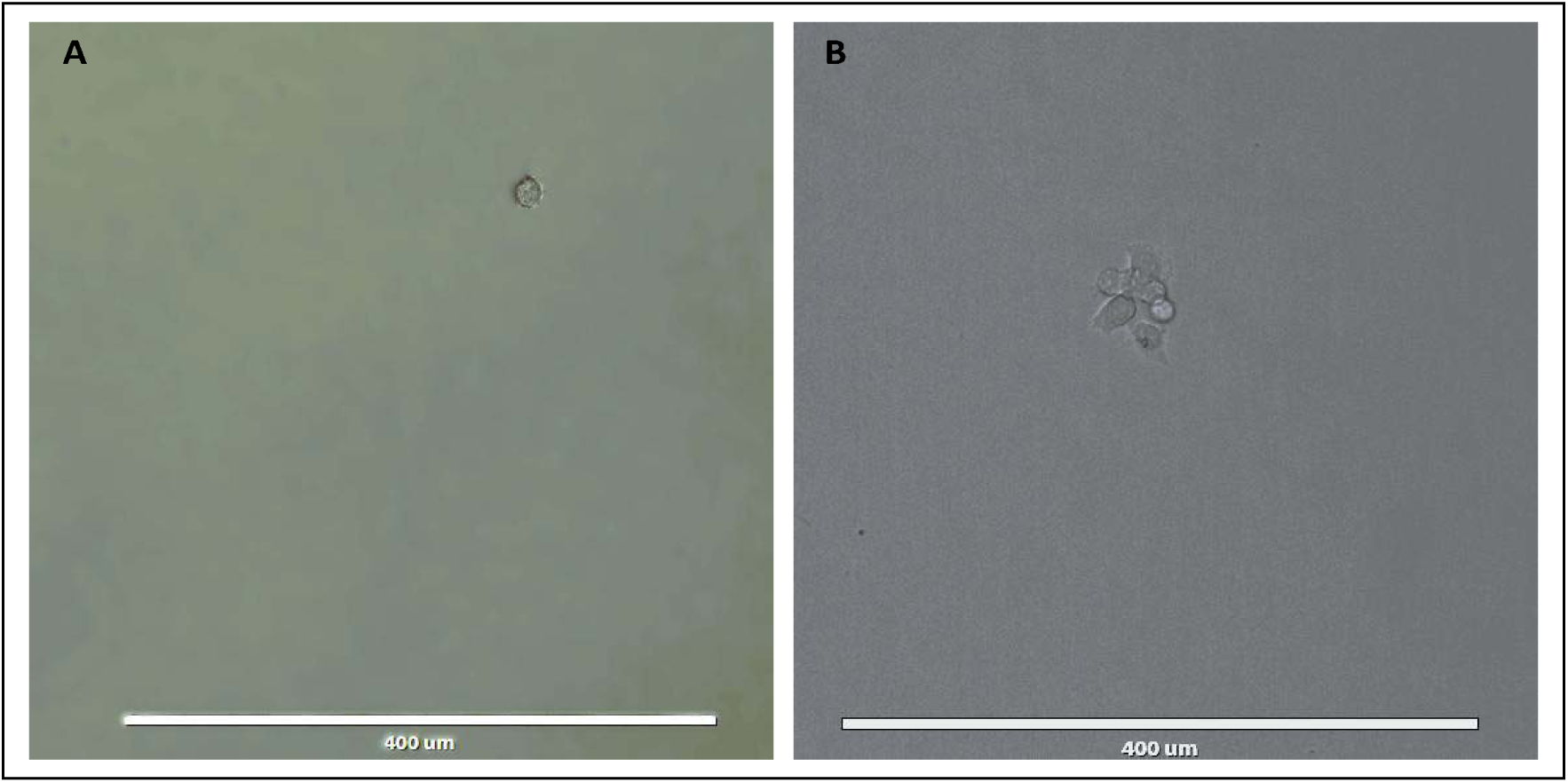
Representative images of (A) single cancer cells or (B) cancer cell clusters. Imaging was performed using an inverted (EVOS) microscope (at10X), and scale bar represents 400 µm.

### 3.4 Detection and Dispensing of OC Cells and Cell Lines

Negative and positive controls established fluorescence gating thresholds (Figures S1, S2). Signal-to-background ratio for FITC staining was 12.4 ± 2.1 (positive event MFI divided by autofluorescence). Low event counts were observed for patient-derived OC samples (Figures S3-S5). For OCAS12, only 15 events were detected after >5 minutes. In contrast, improved detection was achieved for SiHa cells by lowering FSC trigger to 2000 (Figure S6), with 9/10 droplets containing single cells.

### 3.5 Single-Cell Dispensing from Patient and Cell Line Samples

Single-cell dispensing succeeded for all samples (Figures S7-S12). Event rates ranged from 0.5 to 5 events per second, requiring sample-specific FSC optimisation. OCAS12 showed ∼4 events/second (FSC 2000); OCAS14 and OCAST16 showed ∼1.5 and 0.5 events/second (FSC 7000). SiHa cells showed ∼5 events/second (FSC 7000); C33a and HeLa showed ∼1.5 events/second. Coefficient of variation across three runs was 18% for patient-derived samples vs. 12% for cell lines, indicating greater primary sample variability and underscoring the need for pre-sort optimisation.

### 3.6 Dispensing of Cancer Cells Spiked into Blood (Mimicking CTCs)

For Sample 1 (SiHa cell line), a dispensing density of approximately 1.5 events per second was observed with an FSC trigger of 2000. Only FITC+ cells (majority PE−) were selected. A total of 50 target cells were dispensed into two wells, with fewer than 50 cells dispensed into a third well before cartridge depletion (Figure S13). Sample 2 (Hela cell line), analysed under identical conditions, yielded 50 target cells distributed across four wells, with a partial fifth well (Figure S14). Sample 3 (C33a cell line), exhibited a lower dispensing density of approximately 1 event per second, enabling 50 target cells to be dispensed into twelve wells (Figure S15). Sample 4 (OCAS14), with a density of approximately 3 events per second, showed a substantial proportion of dispensed events that were also PE+; nevertheless, 50 target cells were dispensed into ten wells (Figure S16). Sample 5 (OCAS12), displayed a similar density and PE+ profile, resulting in 50 target cells dispensed into nine wells (Figure S17). Following refinement of the isolation strategy, new spiked blood samples were dispensed using an FITC+/PE− gating strategy. The negative control (unstained HeLa cells) was analysed at a density of approximately 4 events per second with an FSC trigger of 5000 (Figure S18).

For spiked blood Sample 1 (C33a cells), a density of approximately 1 event per second was observed with a fluorescent gate set to FITC 1000 and an FSC trigger of 5000 (Figure S19). Sample 2 (OCAS12) was analysed under identical parameters (Figure S20). Sample 3 (HeLa cells) required a higher fluorescent gate (FITC 10,000) with the same FSC trigger (Figure S21). Samples 4 (SiHa cells) and 5 (OCAS14) were analysed with a fluorescent gate of FITC 5000 and an FSC trigger of 5000, exhibiting densities of approximately 1 and 2 events per second, respectively (Figures S22 and S23).

### 3.7 Cell Viability and Proliferation

To assess cell viability and proliferation status following single cell dispensing (Figure 3), cells were imaged immediately after dispensing and then cultured under the conditions described in section 2.6. As seen in Figure 4, by day 3, there was an increase in cell proliferation and replication compared to day 1 (baseline). All three OC (OCAS12, OCAS14 and OCAST16) samples started from an identical baseline (Day 1 = 1) and reached similar terminal fold changes by Day 21 (∼85–94). However, their proliferative status differed markedly: OCAS12 exhibited the earliest and most pronounced increase between Days 3–7, reflected in the highest mean fold change at Day 7 (22.67 ± 11.01) before growth slowed thereafter; OCAS14 showed a more sustained, gradual increase, with the highest net accumulation after Day 14 culminating in the greatest terminal value (93.67 ± 4.16); and OCAST16 had the slowest early expansion (lowest Day 7 mean, 9.33 ± 2.08), but replication accelerated markedly between Days 7–14 and remained robust through Day 21, nearly matching the others by the endpoint (Table 3).

**Figure 3:**
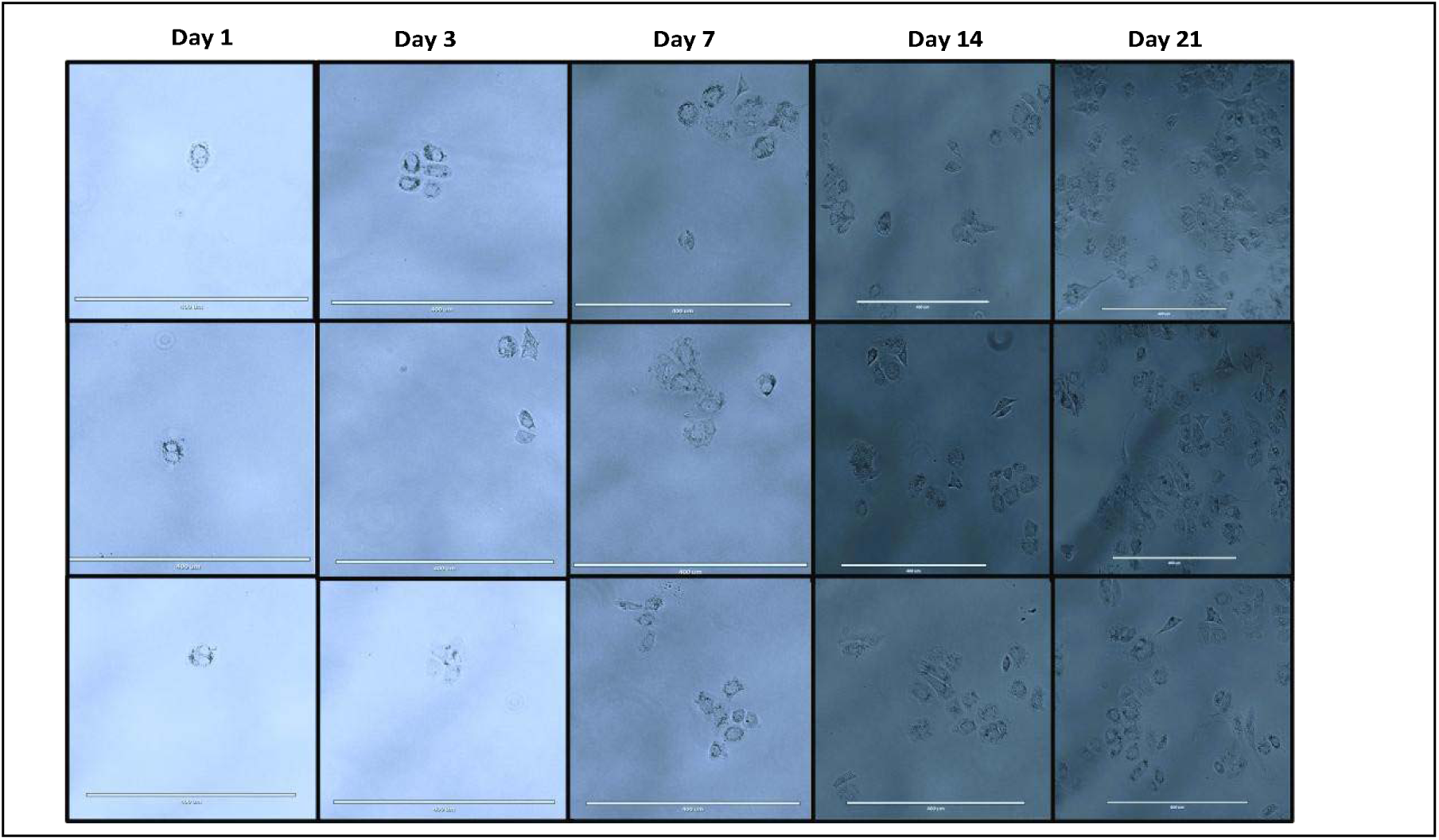
Representative micrographs of single cancer cells following dispensing. Illustrating images of proliferation status over a 21-day culture period. Imaging was performed using an inverted (EVOS) microscope (at 10X), and scale bar represents 400 µm.

**Figure 4:**
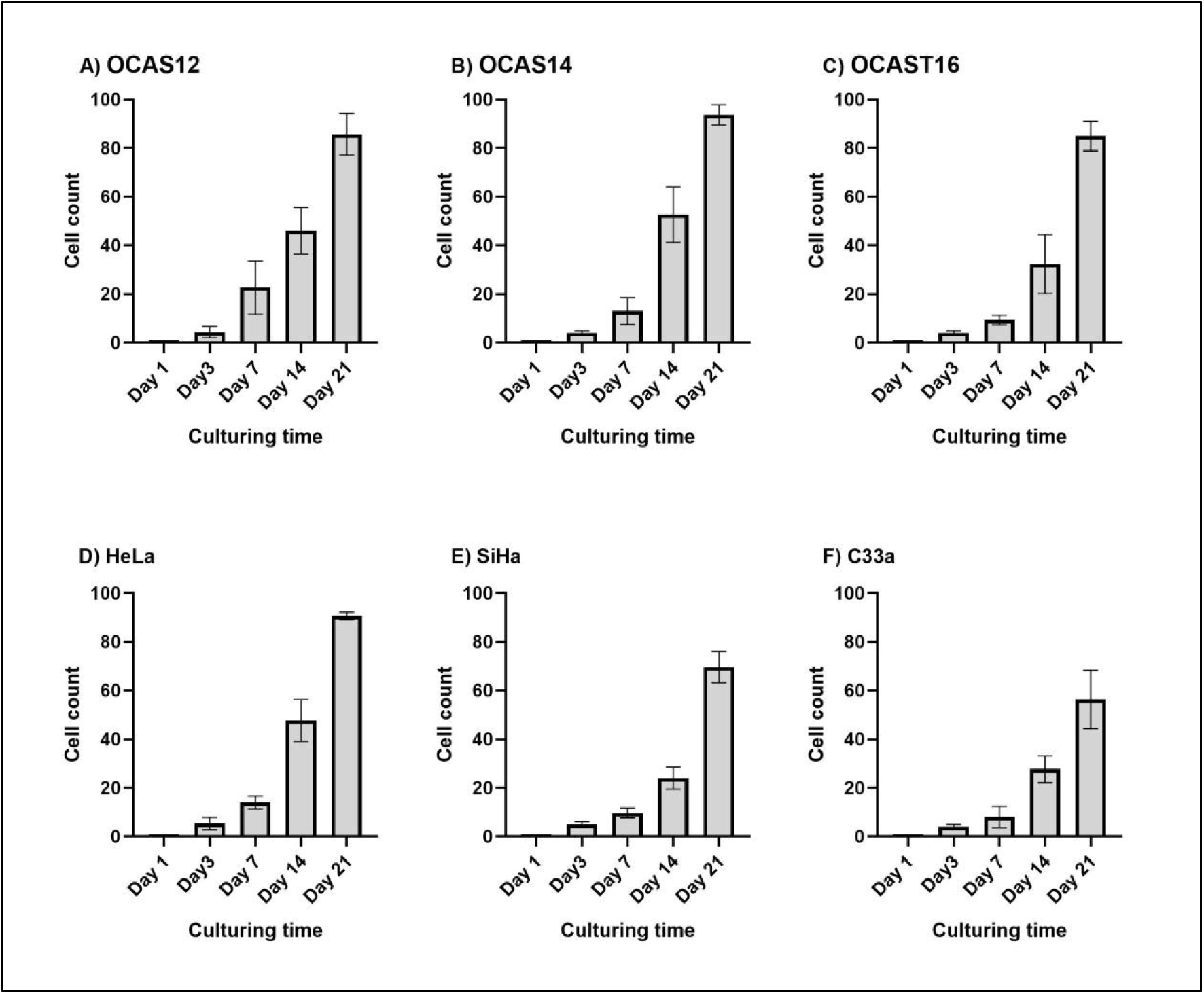
Evaluation of fed-batch culture performance of cell proliferation following Pala™ sorting. The figure depicts the increase in cell number over time in culture (assessed at days 3, 7, 14 and 21), as quantified objectively by three independent investigators.

**Table 3:**
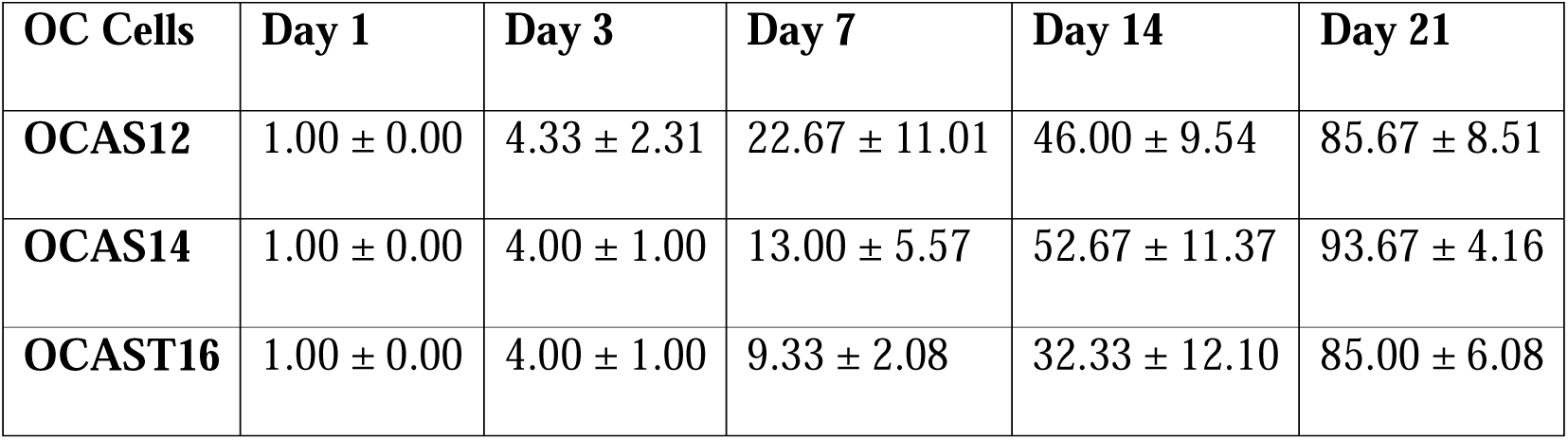
OC Sample Proliferation (Fold Change from Day 1)

| OC Cells | Day 1 | Day 3 | Day 7 | Day 14 | Day 21 |
| --- | --- | --- | --- | --- | --- |
| **OCAS12** | $1.00 \pm 0.00$ | $4.33 \pm 2.31$ | $22.67 \pm 11.01$ | $46.00 \pm 9.54$ | $85.67 \pm 8.51$ |
| **OCAS14** | $1.00 \pm 0.00$ | $4.00 \pm 1.00$ | $13.00 \pm 5.57$ | $52.67 \pm 11.37$ | $93.67 \pm 4.16$ |
| **OCAST16** | $1.00 \pm 0.00$ | $4.00 \pm 1.00$ | $9.33 \pm 2.08$ | $32.33 \pm 12.10$ | $85.00 \pm 6.08$ |

The proliferation rate of all cervical cancer cell lines (HeLa, SiHa, C33a) started from an identical baseline (Day 1 = 1) but diverged substantially in their proliferative kinetics and terminal expansion by Day 21 (Table 4). HeLa cells demonstrated the most robust overall proliferation, reaching the highest mean fold change at Day 21 (90.67 ± 1.53), with a consistent and progressive increase from Day 7 onward (14.00 ± 2.65 at Day 7, 47.67 ± 8.51 at Day 14). In contrast, SiHa cells showed a more moderate but steady expansion, concluding at 69.67 ± 6.43 by Day 21, while C33a cells exhibited the slowest proliferation, with the lowest Day 7 value (8.00 ± 4.36) and the greatest inter-replicate variability at the endpoint (56.33 ± 12.06).

**Table 4:**
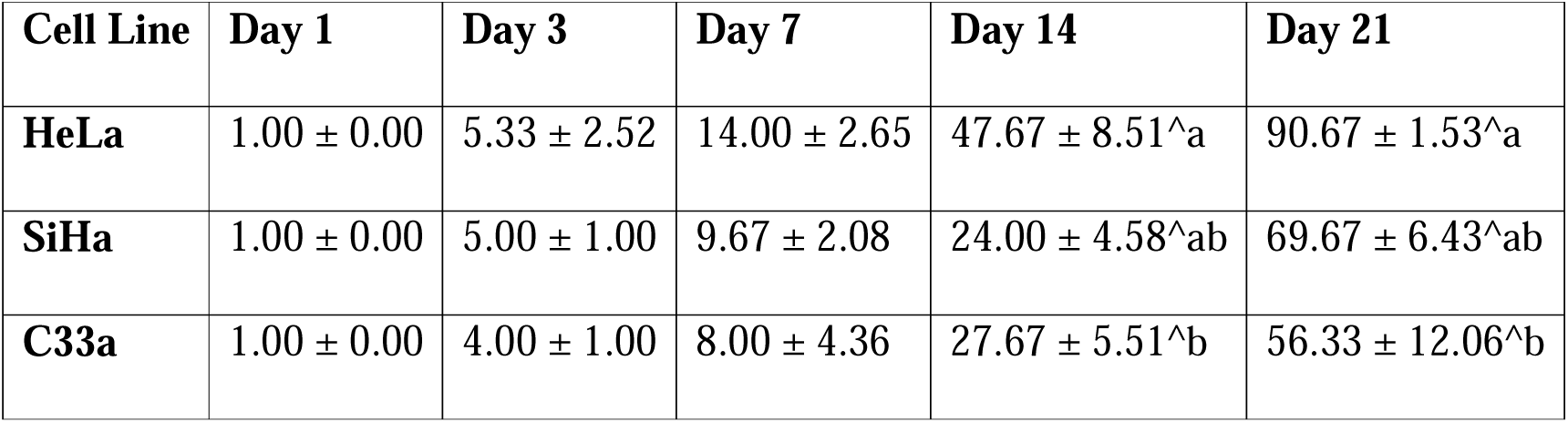
Cervical Cancer Cell Line Proliferation (Fold Change from Day 1)

| Cell Line | Day 1 | Day 3 | Day 7 | Day 14 | Day 21 |
| --- | --- | --- | --- | --- | --- |
| **HeLa** | $1.00 \pm 0.00$ | $5.33 \pm 2.52$ | $14.00 \pm 2.65$ | $47.67 \pm 8.51^{\wedge a}$ | $90.67 \pm 1.53^{\wedge a}$ |
| **SiHa** | $1.00 \pm 0.00$ | $5.00 \pm 1.00$ | $9.67 \pm 2.08$ | $24.00 \pm 4.58^{\wedge ab}$ | $69.67 \pm 6.43^{\wedge ab}$ |
| **C33a** | $1.00 \pm 0.00$ | $4.00 \pm 1.00$ | $8.00 \pm 4.36$ | $27.67 \pm 5.51^{\wedge b}$ | $56.33 \pm 12.06^{\wedge b}$ |

Overall, OC samples as a group achieved uniformly high terminal expansion but differed in the timing of peak replication (early, steady, or delayed). Cervical cell lines spanned a wider range of growth, with HeLa cell line matching the OC samples in net growth, while SiHa and C33a cell lines exhibited significantly lower overall proliferative capacity. These differences may reflect distinct underlying oncogenic drivers or tissue-specific growth regulatory mechanisms.

### 3.8 Patient-Derived Organoid Generation

To generate organoids and enable 3D single-cell culture, dispensing events were performed using all three OC patient samples and three cervical cancer cell lines. Each sample was successfully dispensed as single events into 96-well plates. Forward scatter trigger optimisation (FSC = 7000) enabled reliable single-cell detection and dispensing across all sample types. Successful single-cell dispensing was achieved for all OC patient samples included in this study (Figures S24-S26). The dispensing event rate was sample-dependent. For sample OCAST16, a dispensing density of approximately 0.5 events per second was observed using an FSC trigger of 7000. In contrast, samples OCAS12 and OCAS14 exhibited higher dispensing densities of approximately 2.5 events per second under the same FSC trigger setting. Within one week post-dispensing, viable single cancer cells demonstrated outgrowth and formed clusters consistent with early organoid development. Figure 5 shows representative images from organoid samples generated from all patients’ samples (OCAS12, OCAS14 and OCAST16) used in this study, where clusters of cells visible one day post-dispensing had grown into small organoids within a few days, and generation of 3D organoids was obvious after three weeks.

**Figure 5:**
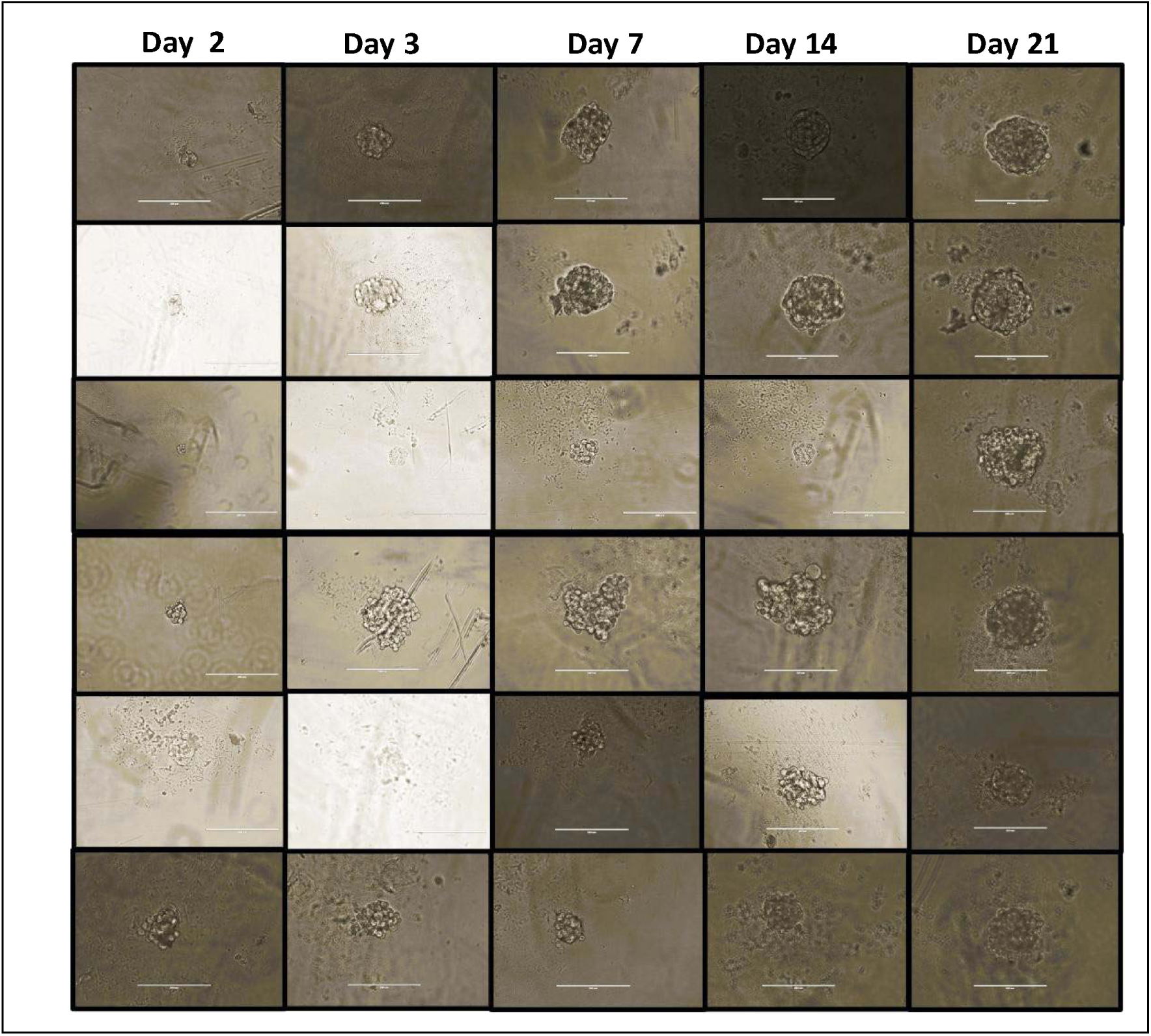
Representative images of organoid samples generated from patient cells and cell lines. Clusters of cells were visible one day after dispensing, and these grew into small organoids within a few days. By three weeks, clear three-dimensional organoid structures had formed. Imaging was performed using an inverted (EVOS) microscope (at 20X), and scale bar represents 200 µm.

### 3.9 Organoid Reconstruction and Marker Expression Profiling

To evaluate whether dissociated organoids (as single cells) could successfully reconstruct three-dimensional PDOs while retaining key phenotypic features, we cultured dispensed single cells from three OC patient samples (OCAS12, OCAS14, OCAST16) under optimised organoid conditions as described in section 2.4.3. Within seven days, viable single cells formed small clusters that progressively developed into organised 3D structures by day 21, confirming successful PDO reconstruction, consistent with the passaging criteria outlined in section 2.4.4.

Following successful reconstruction, we characterised the expression of ten epithelial and cancer associated markers in day 21 PDOs using immunofluorescence and immunohistochemistry (Figure 6), following the staining protocols detailed in section 2.4.4 (lineage marker expression, epithelial and therapeutic marker characterisation). Our phenotypical characterisation results showed that the majority of markers examined were uniformly expressed with consistent localisation patterns. In marked contrast, claudin 5 and MUC5AC showed consistently low to absent expression in all OC PDO lines. To further characterise proliferative activity, we assessed Ki 67 expression. As seen in Figure 7, all three OC PDO lines (OCAS12, OCAS14, OCAST16) showed high levels of Ki 67 expression, indicating active proliferation status.

**Figure 6.**
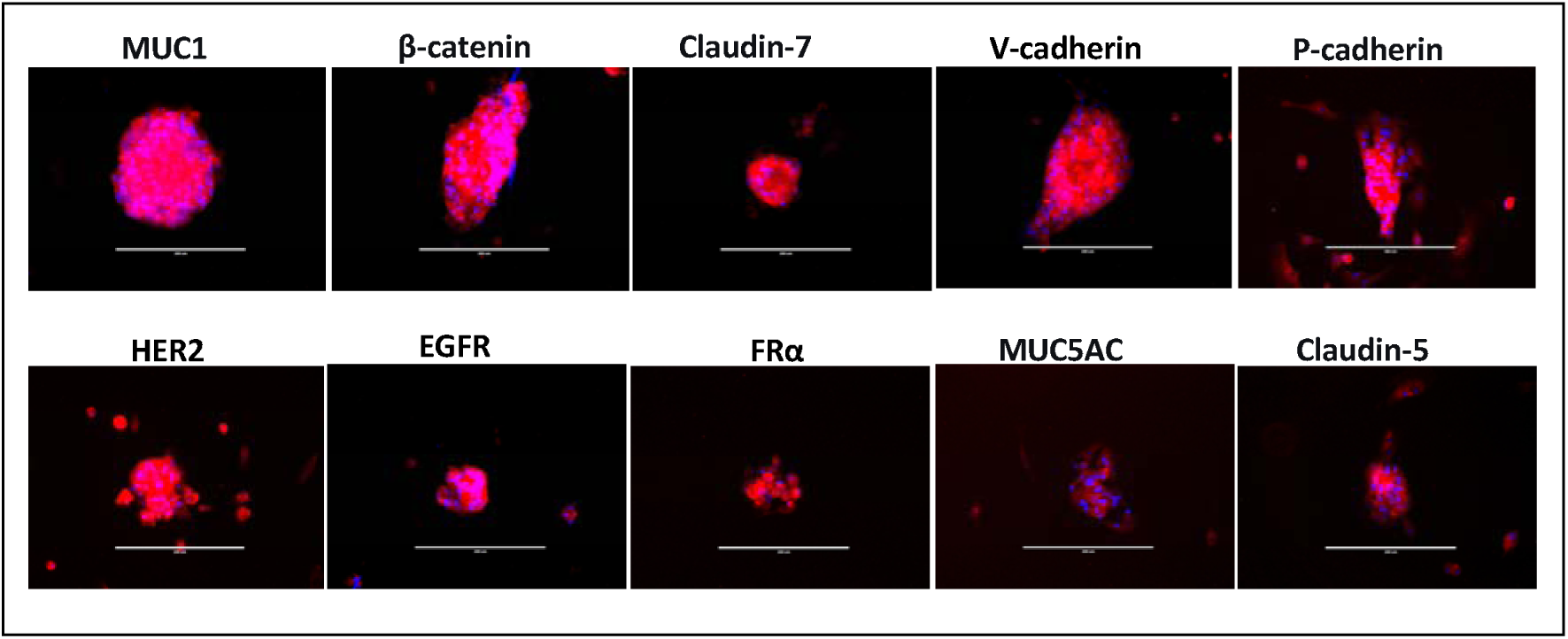
Immunofluorescence and immunohistochemical characterisation of epithelial/cancer markers in day 21 OC PDOs. (MUC1, β-catenin, claudin-7, V-cadherin,P-cadherin, EGFR, FRα, HER2, FRα, MUC5AC and Claudin-5). Imaging was performed using an inverted (EVOS) microscope (at 20X), and scale bar represents 200 µm.

**Figure 7.**
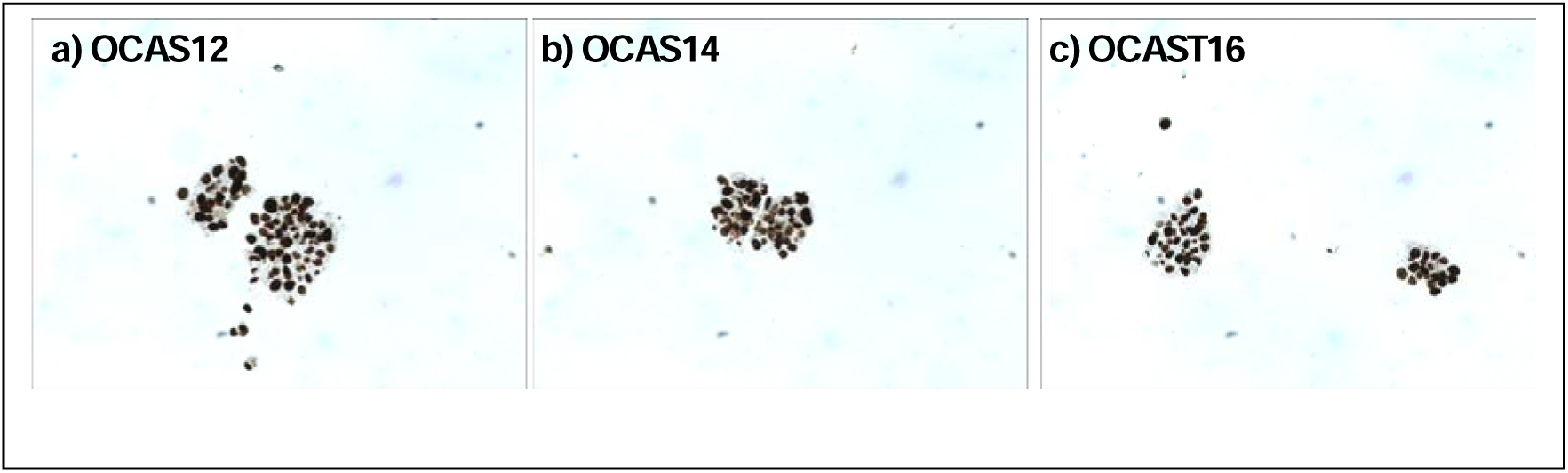
Ki-67 immunoperoxidase staining of OC-PDOs (OCAS12, OCAS14, OCAST16). High nuclear Ki-67 expression indicates active proliferation across all three lines. Imaging was performed using a light microscope at 20X magnification.

## 4. Discussion

The results of this study demonstrate that the Pala™ Single Cell Sorter and Dispenser reliably performs both rare cell isolation from complex blood matrices and gentle single-cell dispensing for organoid generation. Traditional fluorescence activated cell sorting (FACS) instruments operate at pressures between 20 and 70 psi, compromising the viability of sensitive primary cells, and require input cell numbers exceeding 200,000 cells [43,44]. In contrast, the Pala™ operates at <2 psi and processes samples with as few as 100 cells, making it particularly suitable for rare cell applications where every cell is valuable [46,47,50]. These data establish performance benchmarks for internal platform qualification. The demonstrated dispensing efficiency (≥80%) and successful organoid initiation from as few as 100 input cells, together with sustained proliferation over 21 days (85- to 94-fold expansion), define the operating envelope for CTC model isolation and PDO generation workflows.

We propose the following acceptance criteria: (i) single-cell dispensing efficiency ≥70% (≥8/10 droplets containing a single cell); (ii) post-sort viability ≥80% at 24 hours; and (iii) organoid initiation success ≥50% from single cells at day 7. The preservation of cell viability under <2 psi has direct biological implications. High-pressure sorting induces mechanical stress responses, including immediate early gene activation (FOS, JUN), mitochondrial membrane potential disruption, and increased apoptosis within 6–12 hours [43,44]. Our observation of sustained proliferation over 21 days (85- to 94-fold expansion) suggests that Pala™-sorted cells avoid this stress response – particularly critical for primary patient cells that lack the stress adaptation mechanisms of established lines. The Pala™ system addresses limitations inherent to EpCAM-dependent CTC isolation methods such as CellSearch® [12], which miss CTCs undergoing epithelial-mesenchymal transition (EMT) due to EpCAM downregulation [13–15].

Importantly, our use of EpCAM/pan-CK antibodies was for detection, not physical capture. Unlike CellSearch®, which uses EpCAM for magnetic bead-based enrichment, the Pala™ uses fluorescence for optical detection only; cells remain in suspension and are sorted based on user-defined gates without requiring EpCAM expression for physical isolation. A CTC can be detected via cytoplasmic CK even when surface EpCAM is downregulated during EMT. The system’s fluorescence-activated sorting capability allows flexible gating strategies incorporating EMT-associated markers or excluding leukocytes, improving capture of heterogeneous CTC populations [46,47]. We illustrated this by refining our strategy from a broad FITC□ gate (cancer cells plus cancer-immune aggregates) to FITC□/CD45□ for exclusive cancer cell isolation.

The ability to perform both CTC isolation and PDO generation within a single platform enables two clinically relevant scenarios. First, treatment-naïve profiling: CTCs isolated at diagnosis could be expanded as CTC-derived organoids for *ex vivo* drug screening to guide first-line therapy (2–3 week turnaround). Second, resistance monitoring: CTCs collected at progression could identify acquired resistance mechanisms and test alternative regimens. Compared to sequential use of multiple instruments, single-platform workflows reduce sample loss (critical when CTC counts are <10 per 7.5 mL blood) and eliminate inter-instrument variability. Recent microfluidic platforms include Parsortix® PC1 (size-based, FDA-cleared), which recovers viable CTCs but cannot perform marker-based selection, and DEPArray™ (dielectrophoresis), which enables single-cell retrieval but requires pre-enrichment.

The Pala™ uniquely combines fluorescence-activated sorting, low-pressure dispensing, and direct well-plate deposition features not co-existing in any other benchtop system. Compared to FACS, it reduces input requirements 1000-fold (100 vs. 200,000 cells) and maintains higher viability (88% vs. 50–70%) [43,44,46]. Event rates varied substantially across sample types (0.5 to 5 events per second). PDO samples exhibited lower rates (0.5–4 events/second) compared to SiHa cell lines (5 events/second), reflecting differences in cell concentration, size, and sample heterogeneity. Primary dissociated tumour samples contain debris, dead cells, and variable numbers of viable single cells, all influencing event detection. For rare cell applications, lower rates (0.5–1.5 events/second) reduce coincidence events, improving sorting purity, whereas higher rates (5 events/second) increase throughput but require careful trigger optimisation. Adjusting the FSC trigger (2000 to 7000) accommodated sample-specific differences.

The low sorting pressure is particularly critical for organoid generation, where successful culture initiation depends on delivering viable, undamaged single cells [42]. The 70% post-dispense well occupancy and observed viable outgrowth confirm retained proliferative capacity. Our demonstration that PDOs can be successfully reconstructed from as few as 100 single cells aligns with the field’s goal of developing reliable, low-input models for functional precision medicine [50]. This efficiency is comparable to recent studies; for example, Thorel et al. reported successful establishment of long-term OC PDOs that closely recapitulate the tumour of origin and clinical response [51], and Sensi et al. achieved a PDO derivation initiation efficiency of 83.3% using optimised protocols [52]. The establishment of stable phenotypes across three independent passages is particularly encouraging, as a major challenge in the field is the long-term stability and faithful expansion of these models [51]. Studies have shown that PDOs can be passaged long-term while maintaining the transcriptomic and genomic profile of their tissue of origin, underscoring their potential for repeated drug testing over extended periods [51].

The demonstration of uniform expression of ovarian epithelial markers including (MUC1, β-catenin, claudin-7, V-cadherin, and P-cadherin) and therapeutic targets (EGFR, FRα, and HER2) across all three OC-PDO lines confirms their epithelial identity and faithful capture of the original tumour’s biomarker landscape. The maintenance of a high proliferation state, indicated by strong Ki-67 expression, is a key functional characteristic of PDOs and is critical for their utility in drug screening [53]. This is consistent with work by Thorel et al., who demonstrated that established ovarian PDO lines can be efficiently used for drug screening, including for PARP inhibitors [51], and with previous findings that Ki-67 and PAX8 expression patterns in organoids mirror those seen in patient tissues, confirming their proliferative state [53].

The markedly lower expression of claudin-5 and MUC5AC in our OC-PDOs, rather than indicating a failure of the model, likely reflects the specific histological subtype of the original tumours and the biological process of organoid culture. The three OC patient samples (OCAS12, OCAS14, OCAST16) were identified as high-grade serous carcinoma (HGSC), and the marker expression is entirely consistent with this pathology. A comprehensive study by Soini et al. demonstrated that claudin-5 is expressed in ovarian epithelial tumours with a “significantly lower frequency than claudins 1, 4, and 7,” making lower expression plausible in epithelial-type models [54]. Claudin-5 is expressed in a subset of HGSCs, often in a microenvironment-dependent manner; the re-cellularisation process into a 3D gel-like matrix, devoid of the original tumour’s vascular niche, could lead to the downregulation of such microenvironment-dependent proteins that may not be essential for cancer cell proliferation in this format [52].

Regarding MUC5AC, its low expression is a powerful validation of our model’s fidelity, not a limitation. Research has established that MUC5AC expression is associated with mucinous and endometrioid ovarian carcinomas, whereas high-grade serous carcinomas are only “rarely MUC5AC positive,” with a prevalence of approximately 5% [55]. Therefore, the low to absent expression of MUC5AC in our PDOs accurately reflects their HGSC origin and is consistent with a large-scale analysis of MUC5AC expression across 603 ovarian cancers [55]. The low or absent expression of this mucin is thus a marker of a HGSC phenotype and suggests a potentially more aggressive tumour biology. Collectively, these findings establish that the Pala™ platform not only enables successful single-cell dispensing and organoid reconstruction but also preserves the molecular and proliferative characteristics of the original patient tumours. The distinct expression patterns of MUC5AC and claudin-5 serve as robust internal controls, validating that the model has correctly retained its tumour-of-origin characteristics.

While spiked cell line experiments provide reproducible benchmarking, they cannot replicate the biological complexity of authentic patient CTCs, which exhibit greater fragility, phenotypic plasticity, and leukocyte association [4]. This study establishes technical feasibility but not clinical validation. Future studies using prospectively collected CTC samples from OC patients (n ≥ 10) should validate recovery rates (target: ≥70%), viability (target: ≥80%), and CTC-PDOs generation success (target: ≥50%). Similarly, long-term stability (>3 passages) and serial passaging capacity were not assessed, as our primary objective was establishing organoid initiation and the ability of re-initiation post-dissociation a prerequisite step where many platforms fail. As the pharmaceutical industry increasingly adopts human-relevant preclinical models, technologies enabling PDOs generation and rare circulating cell isolation will play essential roles [2,3]. The Pala™ platform, as characterised in this study, provides a versatile, accessible tool for advancing these capabilities within drug discovery pipelines.

## 5. Conclusion

This study demonstrates successful application of the Pala™ Single Cell Dispenser for rare cell isolation from spiked blood samples (mimicking CTCs) and generation of PDOs from dissociated OC samples. The system maintained ≥80% dispensing efficiency across both applications [46,47], while low-pressure operation (<2 psi) preserved cell viability, enabling single-cell outgrowth and early organoid formation [42,46,47,49]. Fluorescence-activated sorting with user-defined parameters enabled flexible isolation of cancer cells, clusters, and cancer-immune aggregates [46–48].

The ability to perform both rare cell isolation and organoid generation within a single platform supports integrated precision oncology workflows bridging liquid biopsy and functional drug testing [3,20–31]. Compared to EpCAM-dependent CTC isolation methods, the fluorescence-activated approach enables flexible multi-marker selection, potentially capturing CTC populations missed by single-marker enrichment strategies [12–15]. While this technical validation used spiked cell lines for controlled benchmarking, independent validation with prospectively collected patient CTC samples (n ≥ 10 per indication) is necessary to establish clinical utility. We provide performance specifications and operating parameters to design such clinical validation studies. The Pala™ platform offers distinct advantages for rare cell applications lower input requirements, reduced sample loss, and gentle sorting conditions that preserve cell viability [43,44,46,47,49] providing a resourceful tool for advancing these capabilities within drug discovery workflows.

## Acknowledgements

We sincerely thank the patients who kindly provided samples for this study. Their generous contribution was essential to this research. We also thank Bio-Techne for providing the Pala™ Single Cell Dispenser and for technical support. This work was supported by the Trinity St James’s Cancer Institute and Eurofins Bioamines.

## Supplemental Information

## Supplementary Results

## 1. Fluorescence Gating Strategy

**Figure S1:**
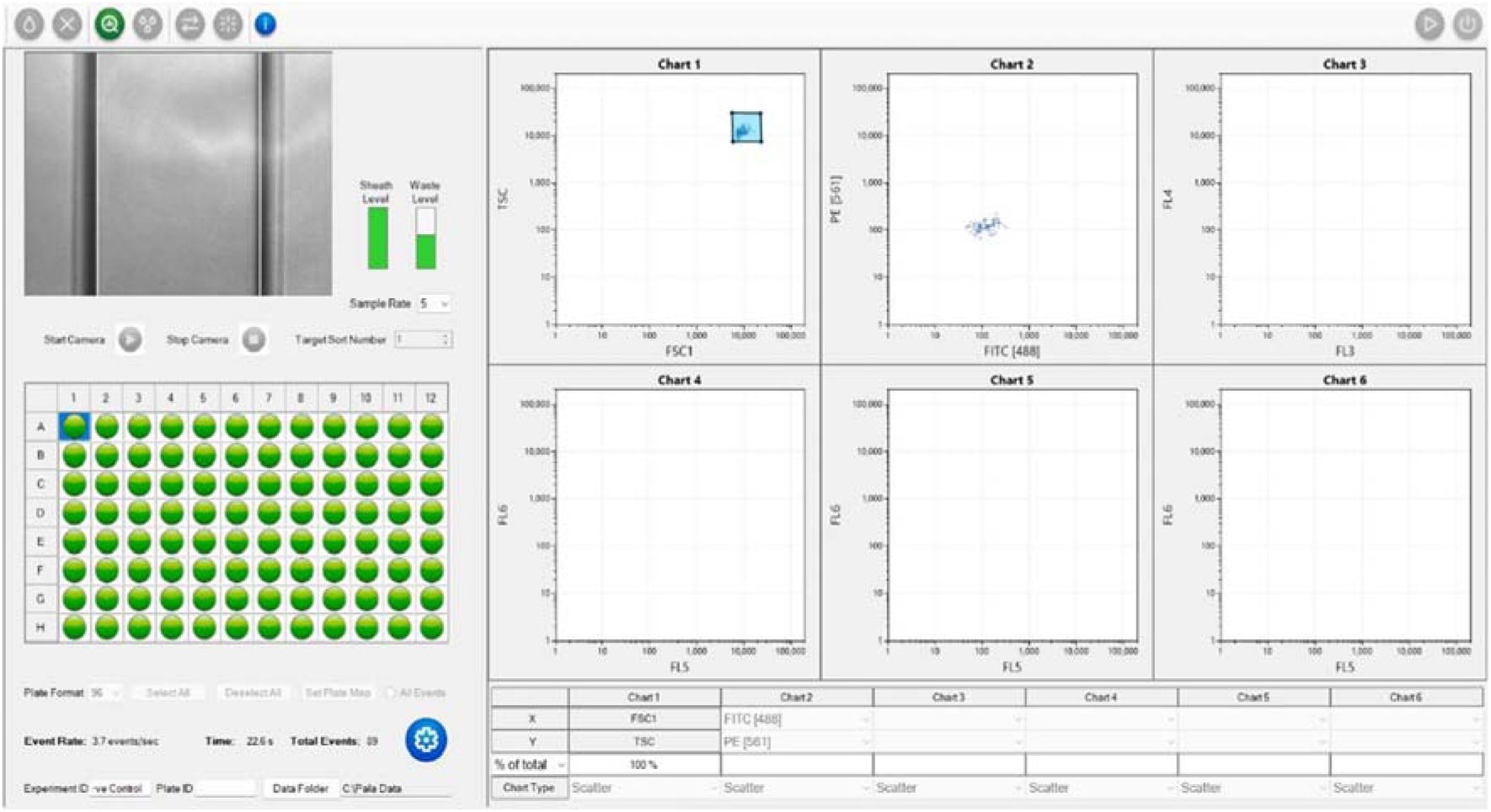
Analysis of the negative control sample (Hela cell line). Sample density was ∼3 events/sec, with the FSC trigger set to 5000. Analysis shows that events >1000 in the FITC and PE channels can be considered positive events.

**Figure S2:**
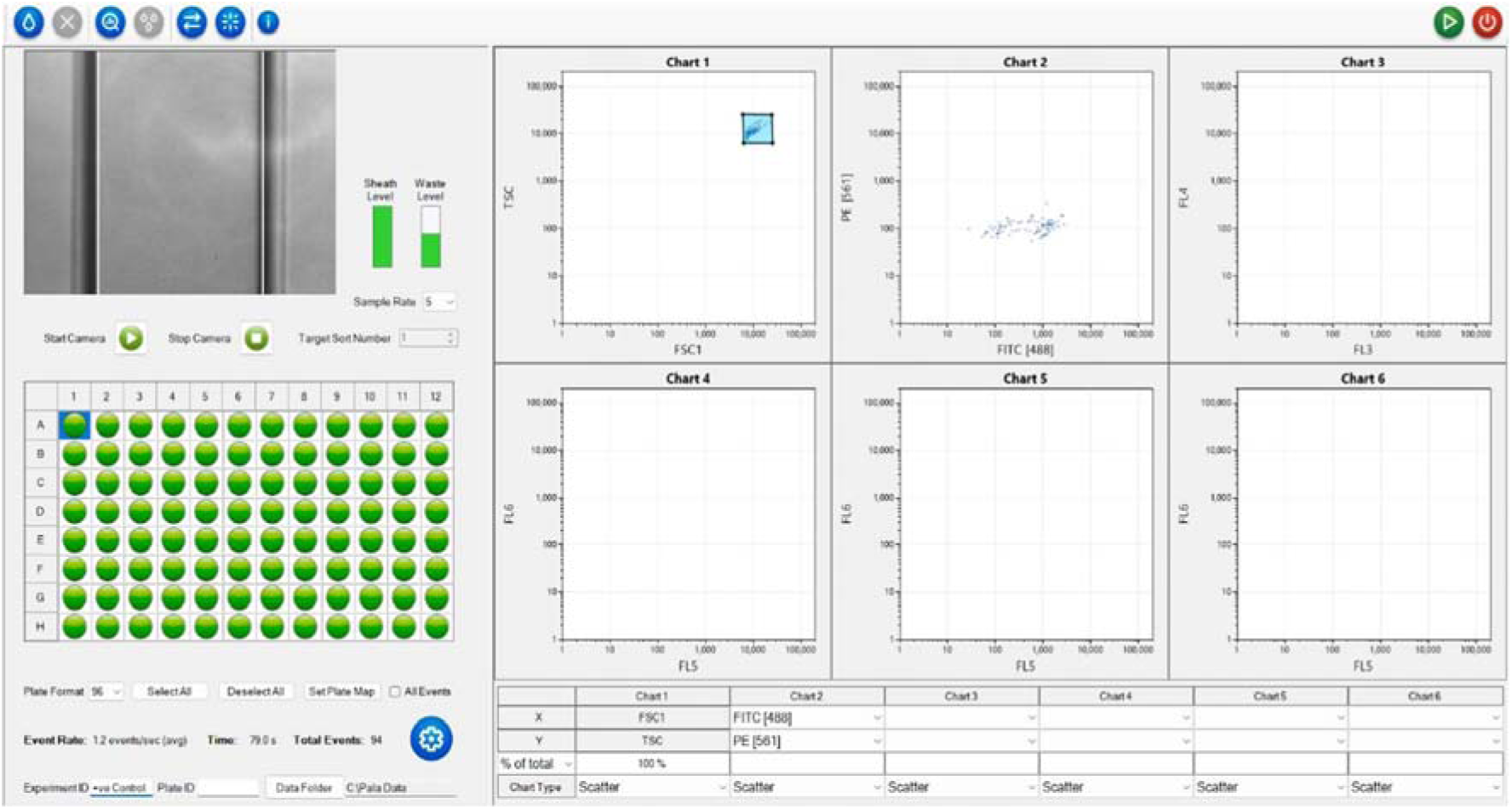
Analysis of the positive control sample (Hela cell line). Sample density was ∼3 events/sec, with the FSC trigger set to 5000. Analysis shows that not all events were FITC+, but a significant population of cells populated the chart at >1000 in the FITC channel. Subsequent gates were drawn to isolate events that populate at >1000 in the FITC channel.

**Figure S3:**
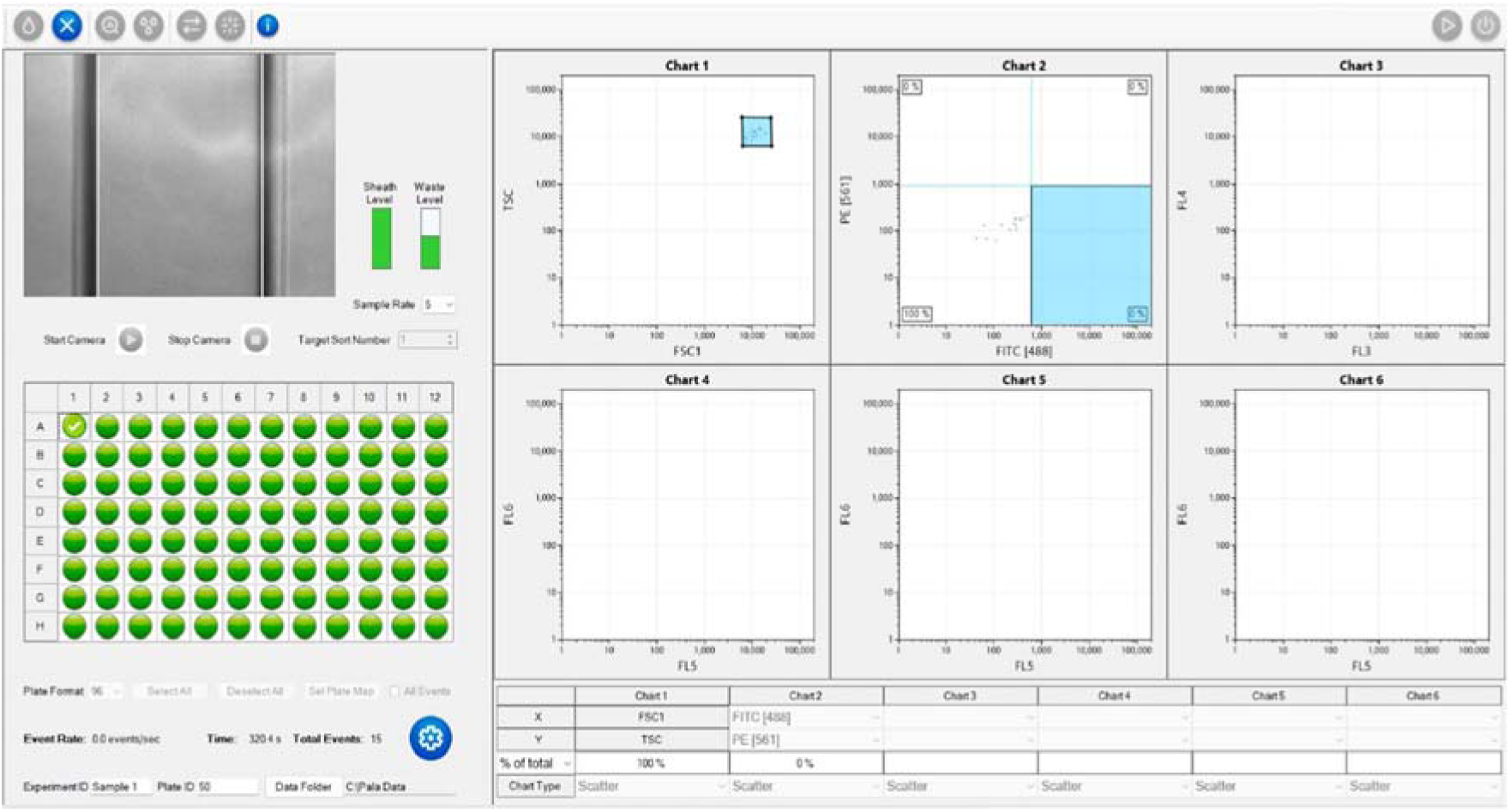
Analysis of Sample 1 (OCAS12). Only 15 events were detected after >5 minutes of analysis, indicating that most of the sample had been lost during processing.

**Figure S4:**
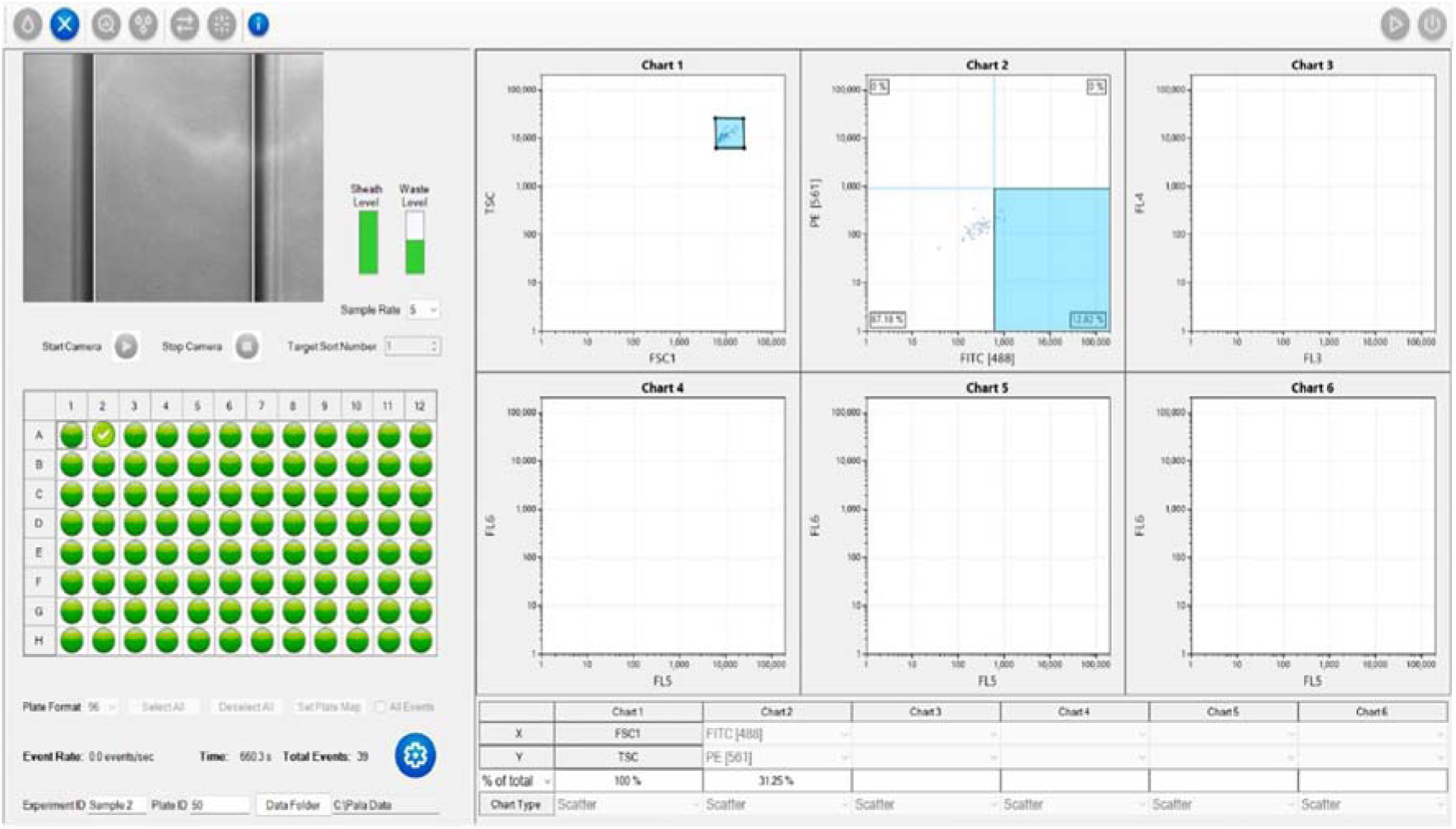
Analysis of Sample 2 (OCAS14). Low cell counts were detected, but some target cells were present.

**Figure S5:**
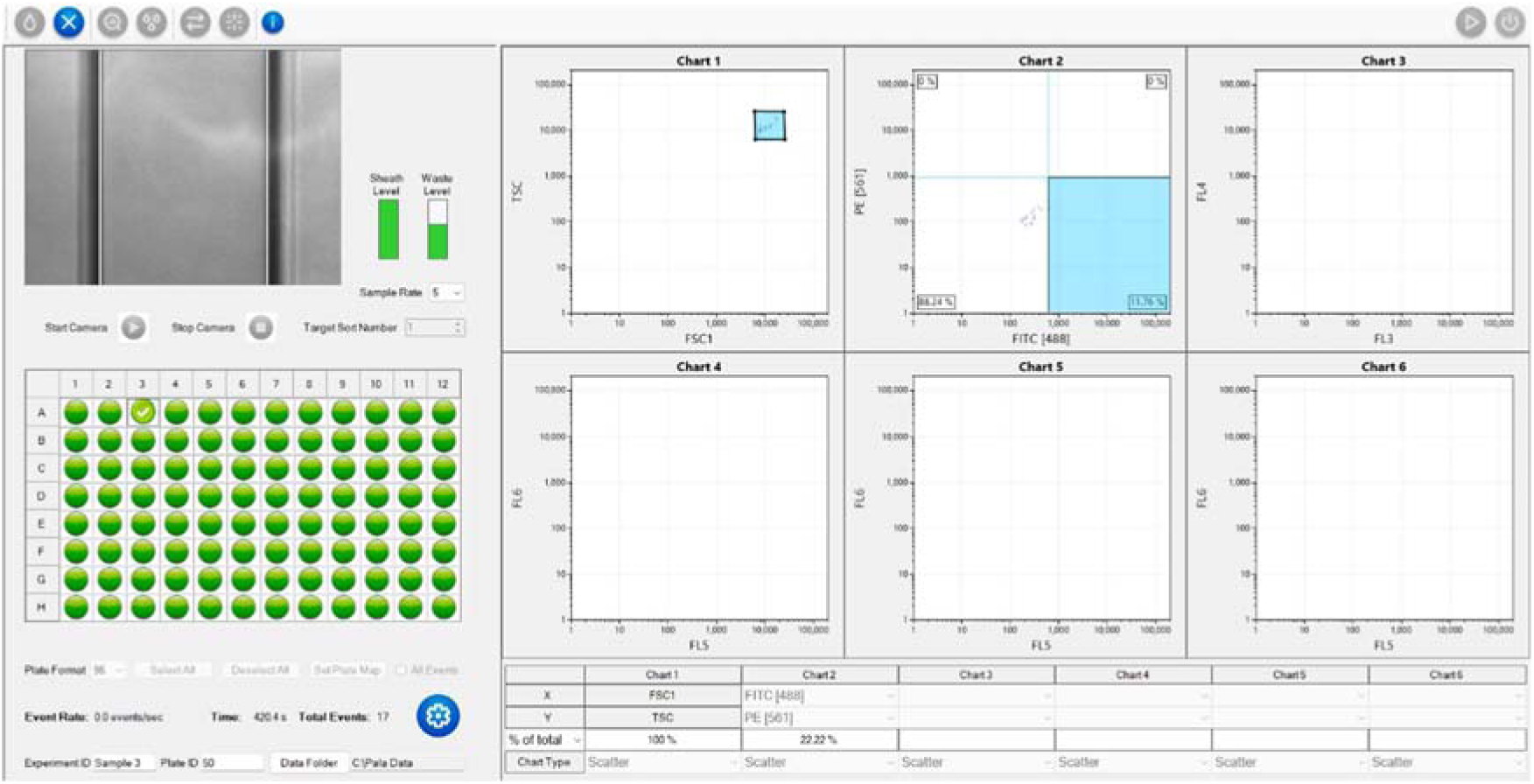
Analysis of Sample 3 (OCATS16). Similar low cell counts were detected as in samples 1 and 2, but some target cells were detected at low fluorescence.

**Figure S6:**
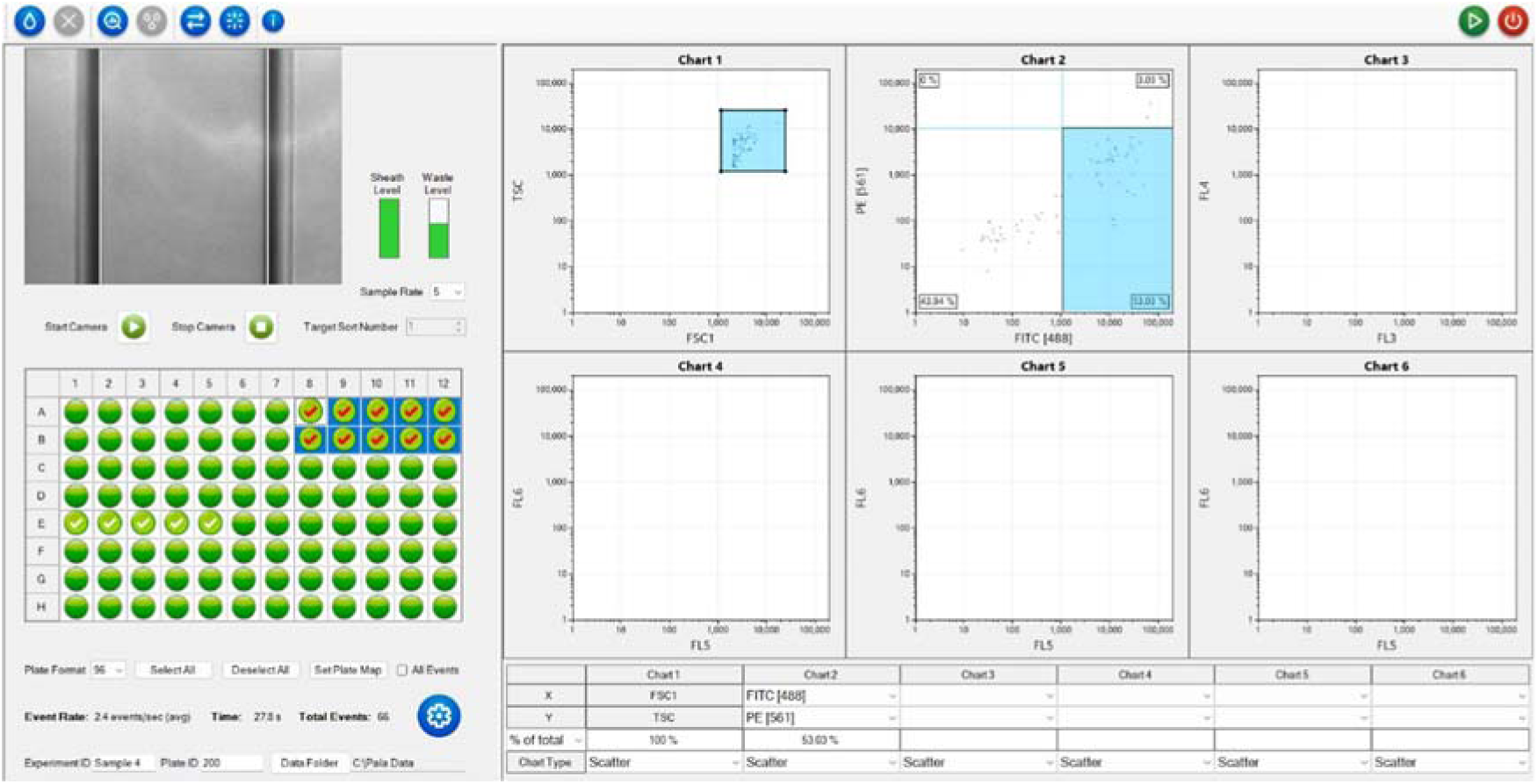
Analysis of Sample 4 (SiHa cell line). FSC trigger level lowered to 2000, higher cell counts observed, and higher numbers of target cells detected at high fluorescence.

## 2. Single-Cell Dispensing from Patient and Cell Line Samples

**Figure S7:**
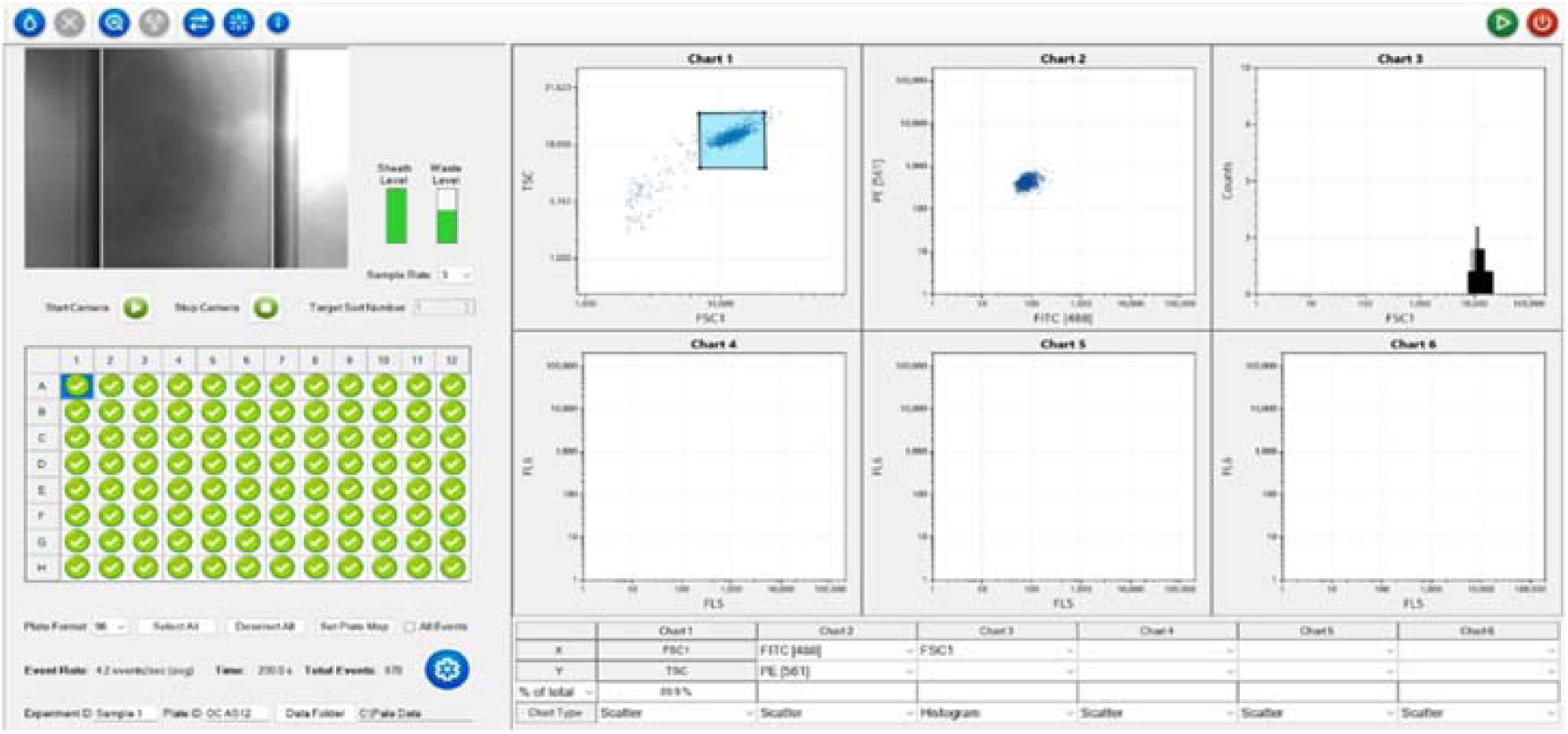
Single cell dispenses of Sample 1 (OC AS12). Sample density was ∼4 events/sec, with an FSC trigger set to 2000.

**Figure S8:**
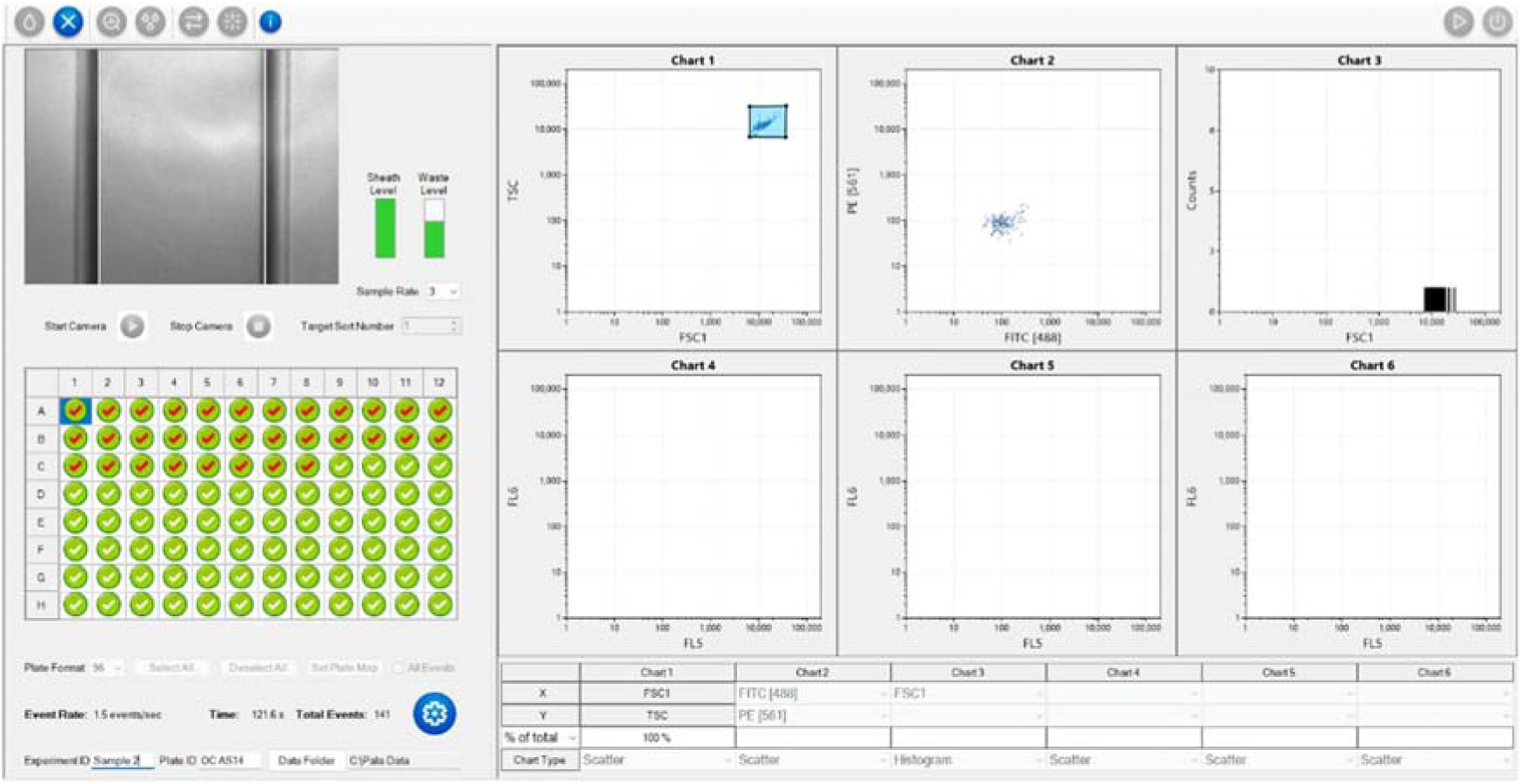
Single cell dispenses of Sample 2 (OC AS14). Sample density was ∼1.5 events/sec, with an FSC trigger set to 7000.

**Figure S9:**
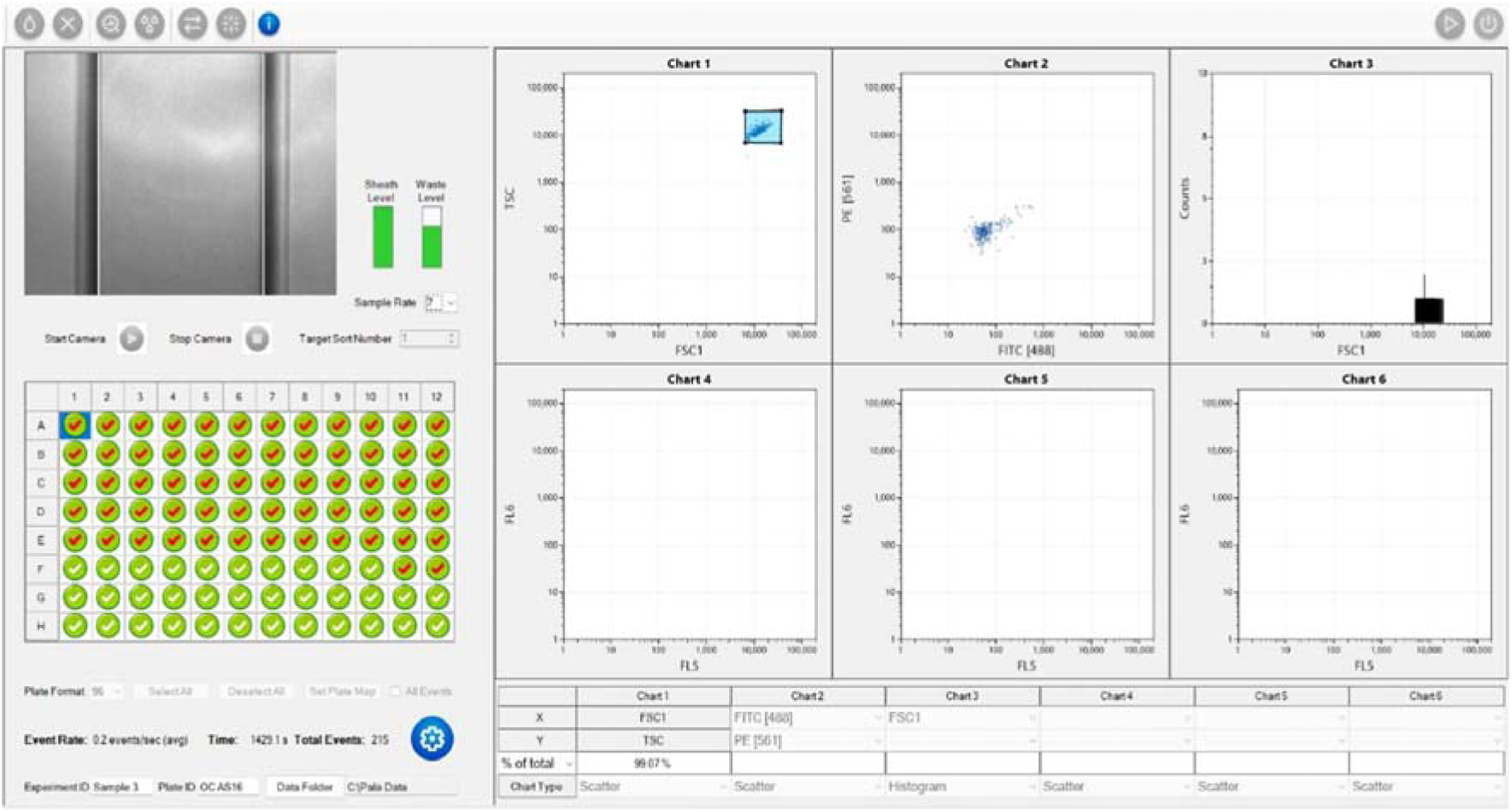
Single cell dispenses of Sample 3 (OC AST16). Sample density was ∼0.5 events/sec, with an FSC trigger set to 7000.

**Figure S10:**
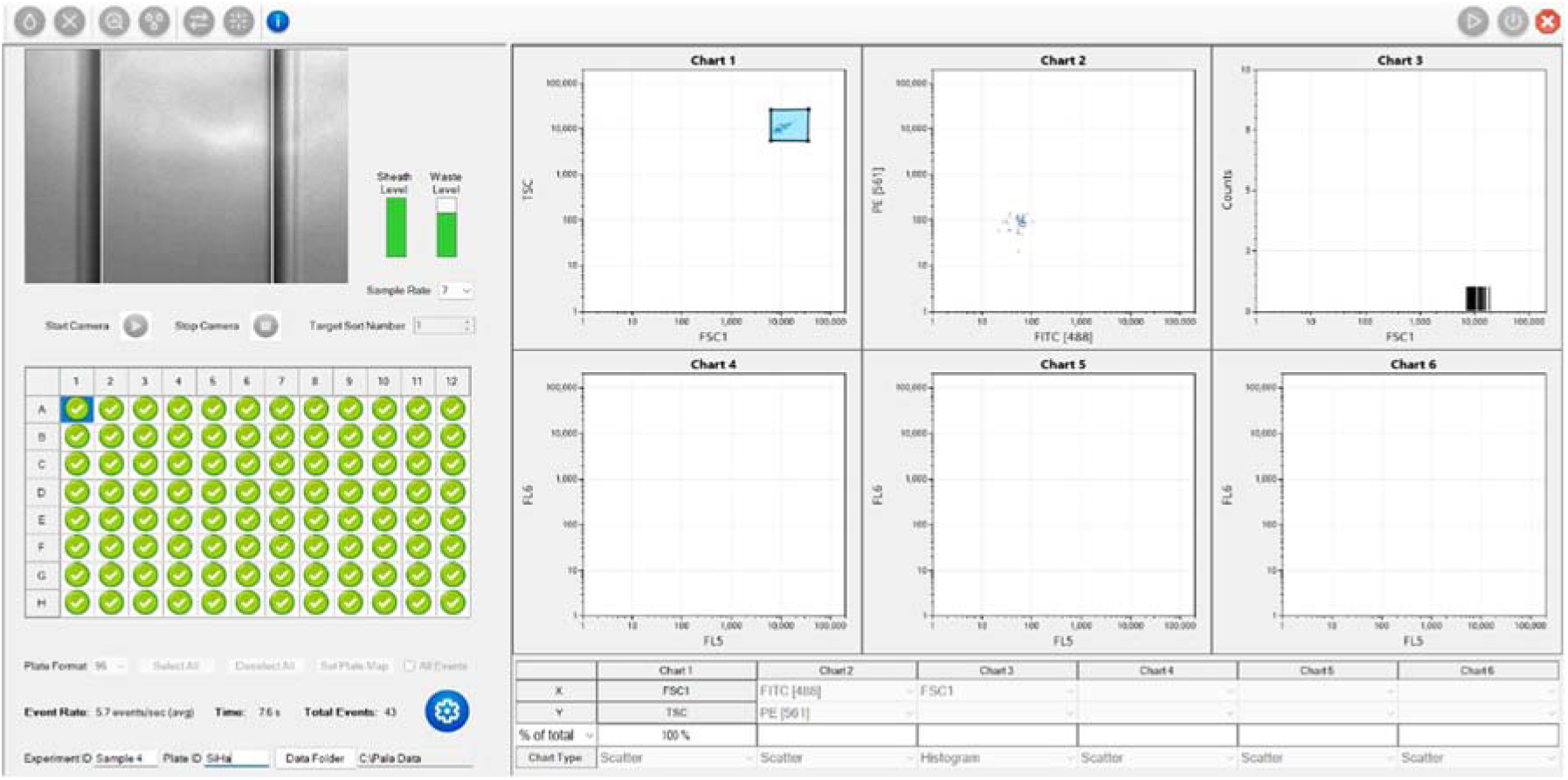
Single cell dispenses of Sample 4 (SiHa cell line). Sample density was ∼5 events/sec, with an FSC trigger set to 7000.

**Figure S11:**
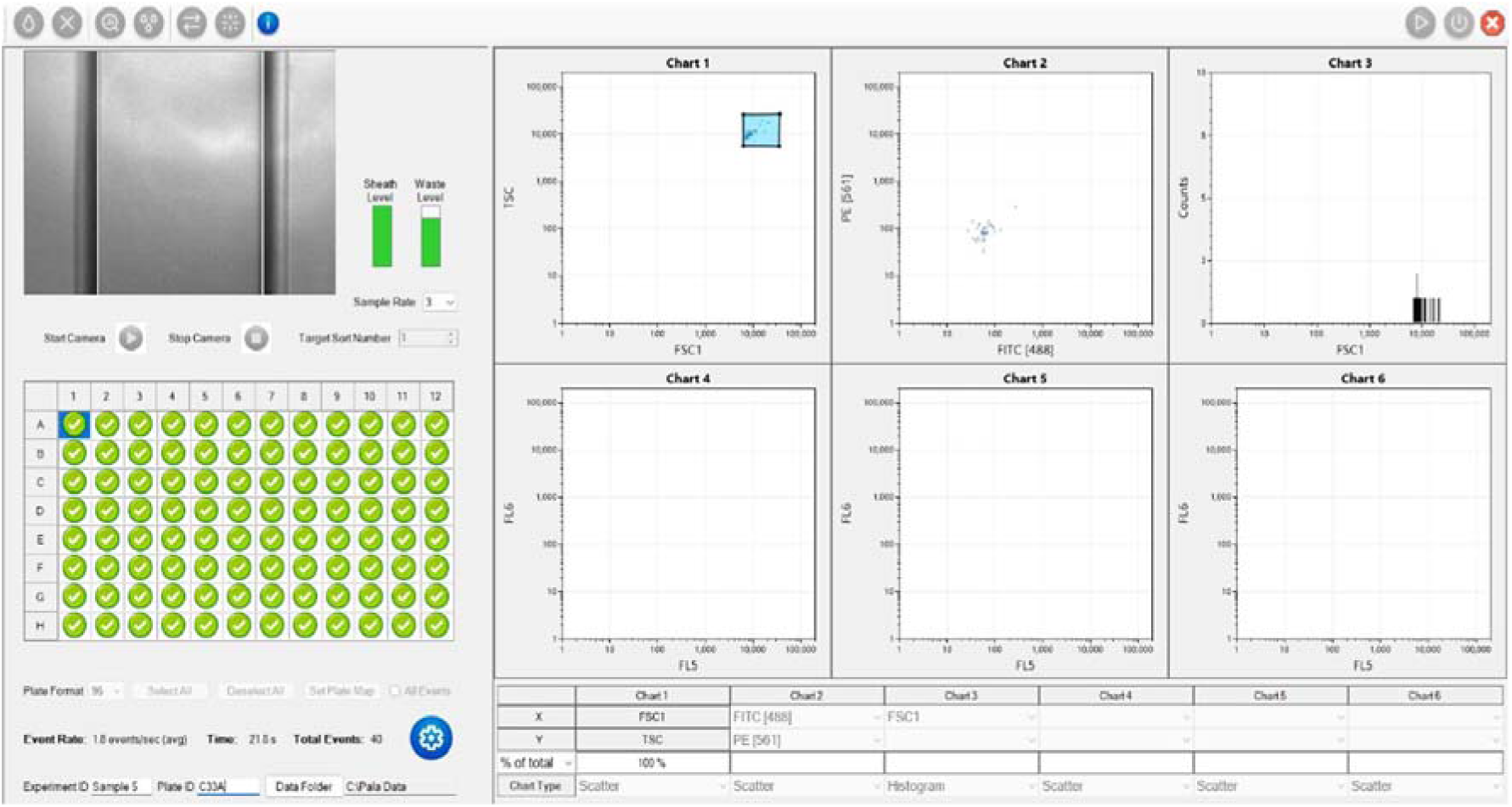
Single cell dispenses of Sample 5 (C339). Sample density was ∼1.5 events/sec, with an FSC trigger set to 7000.

**Figure S12:**
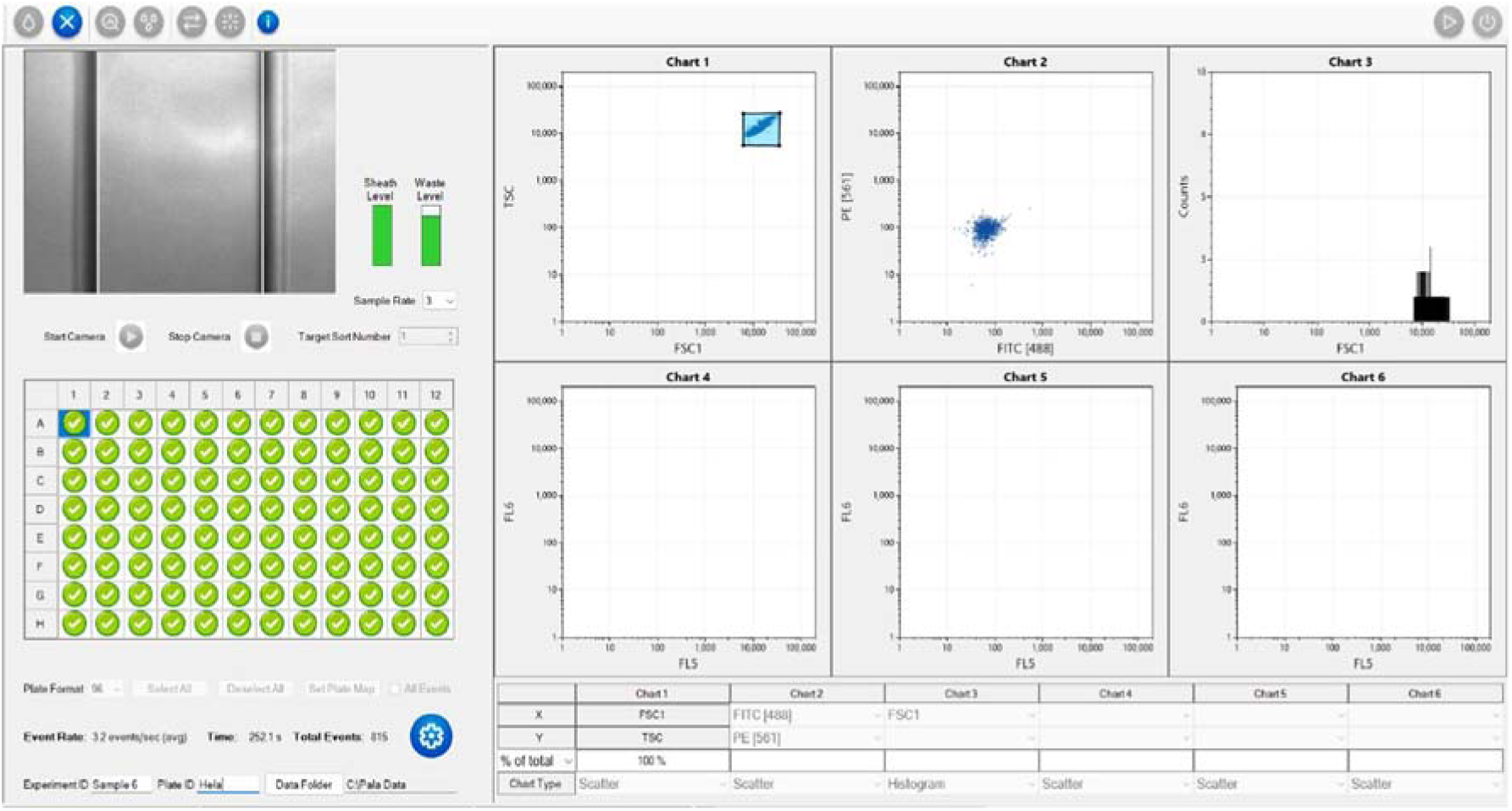
Single cell dispenses of Sample 6 (HeLa cell line). Sample density was ∼1.5 events/sec, with an FSC trigger set to 7000.

## 3. Cancer Cells Spiked into Blood (mimicking CTCs) Sample

### a. Optimisation of CTC cells dispensing

**Figure S13:**
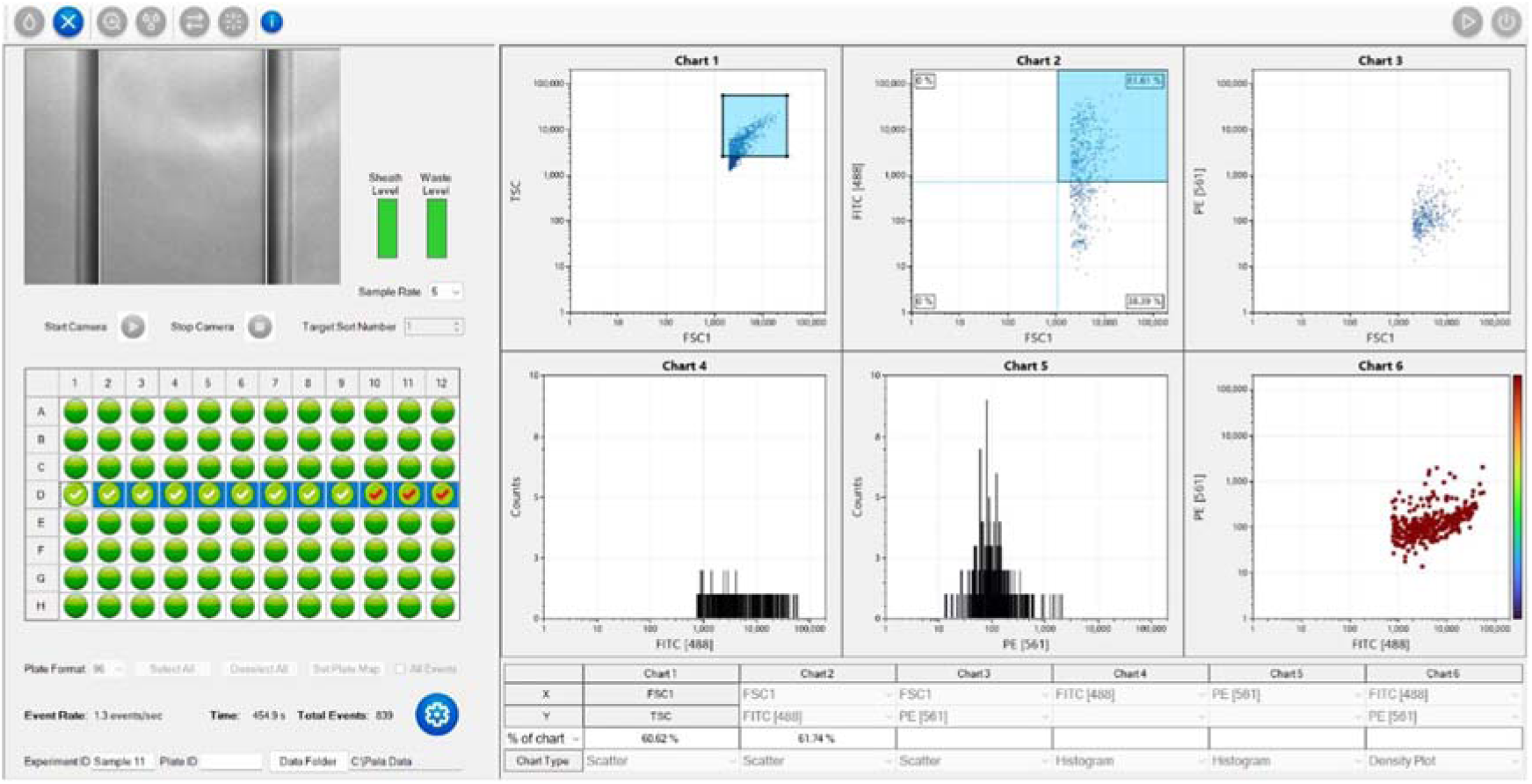
Single cell dispensing of Sample 1 (SiHa cell line). Sample density was ∼1.5 events/sec with the FSC trigger level set to 2000. FITC+ only were selected, with the majority observed as PE−. Fifty target cells were dispensed into 2 wells, with <50 cells dispensed into a third well before the cartridge emptied.

**Figure S14:**
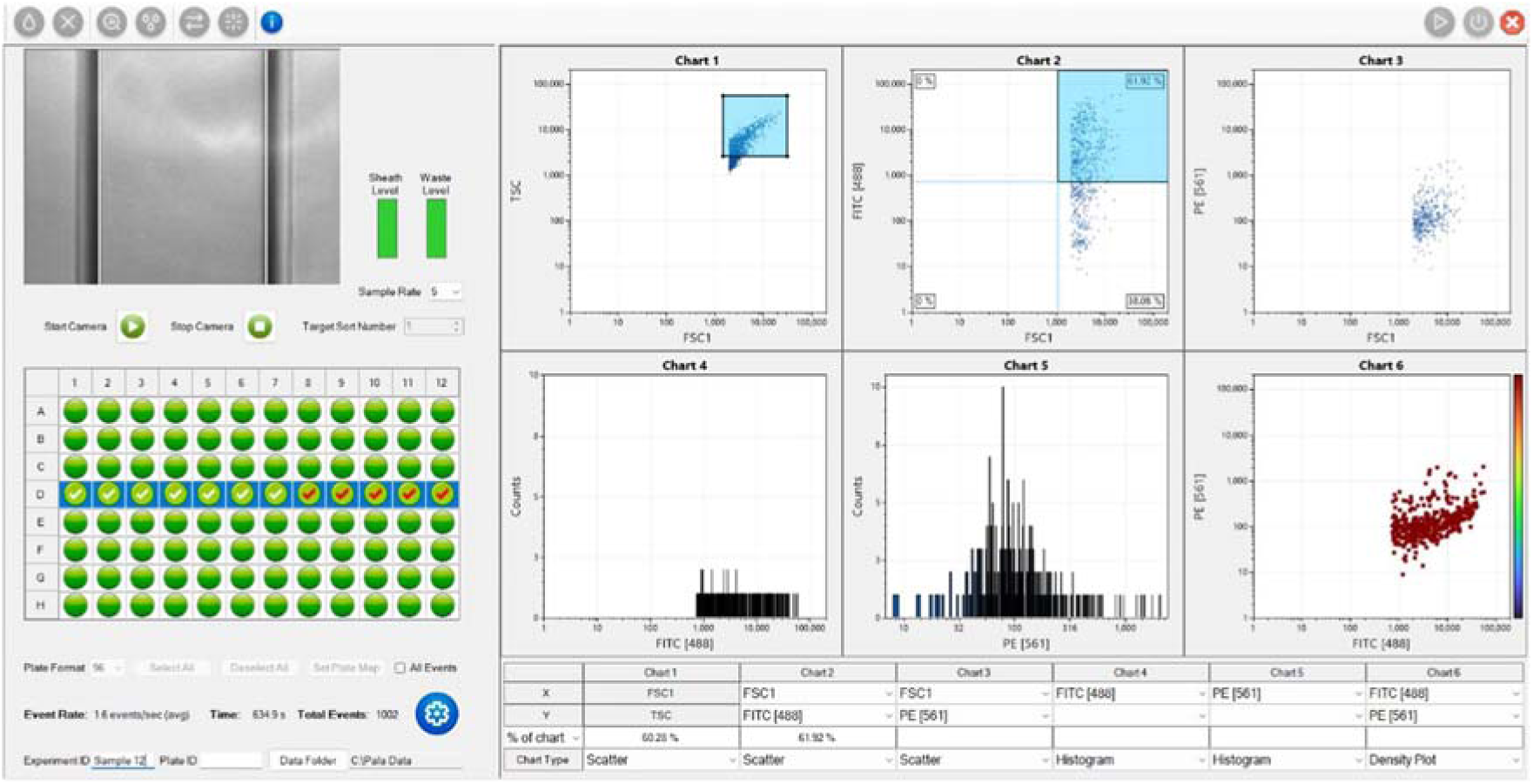
Single cell dispensing of Sample 2 (Hela cell line). Sample density was ∼1.5 events/sec with the FSC trigger level set to 2000. FITC+ only were selected, with the majority observed as PE−. Fifty target cells were dispensed into 4 wells, with <50 cells dispensed into a fifth well before the cartridge emptied.

**Figure S15:**
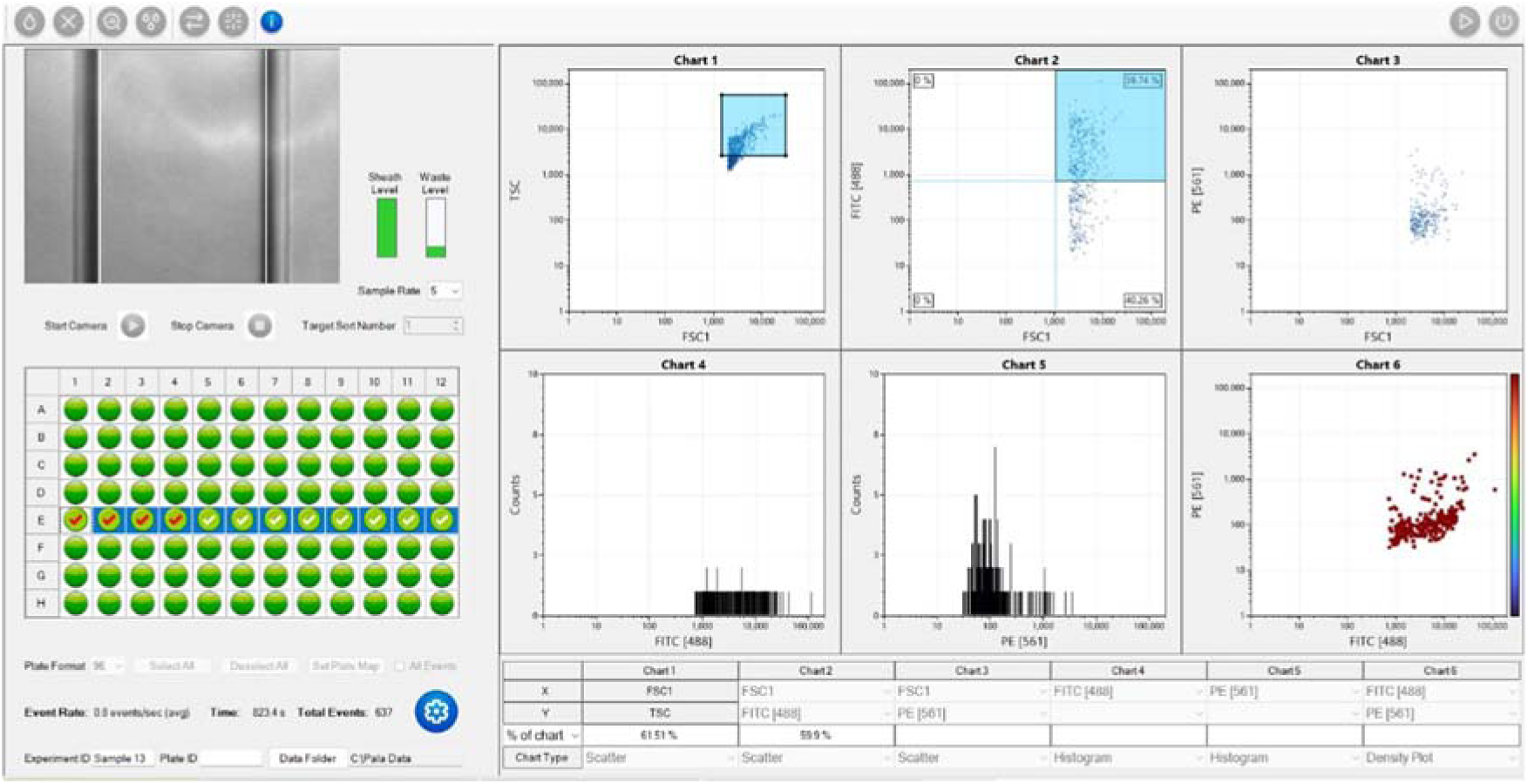
Single cell dispensing of Sample 3 (C33a cell line). Sample density was ∼1 event/sec with the FSC trigger level set to 2000. FITC+ only were selected, with the majority observed as PE−. Fifty target cells were dispensed into 12 wells.

**Figure S16:**
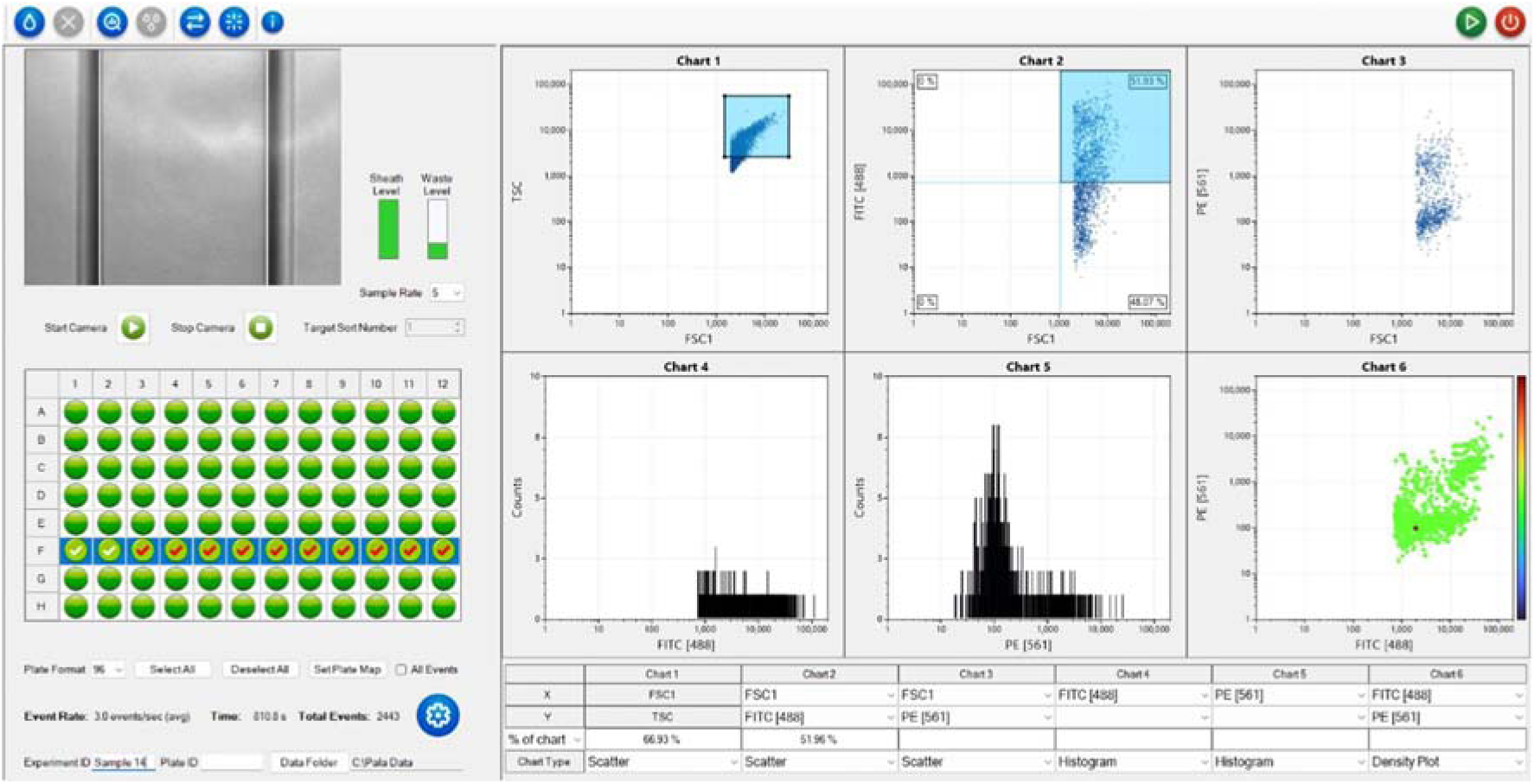
Single cell dispensing of Sample 4 (OCAS14). Sample density was ∼3 events/sec with the FSC trigger level set to 2000. FITC+ only were selected, with a large number of dispensed events observed as also being PE+. Fifty target cells were dispensed into 10 wells.

**Figure S17:**
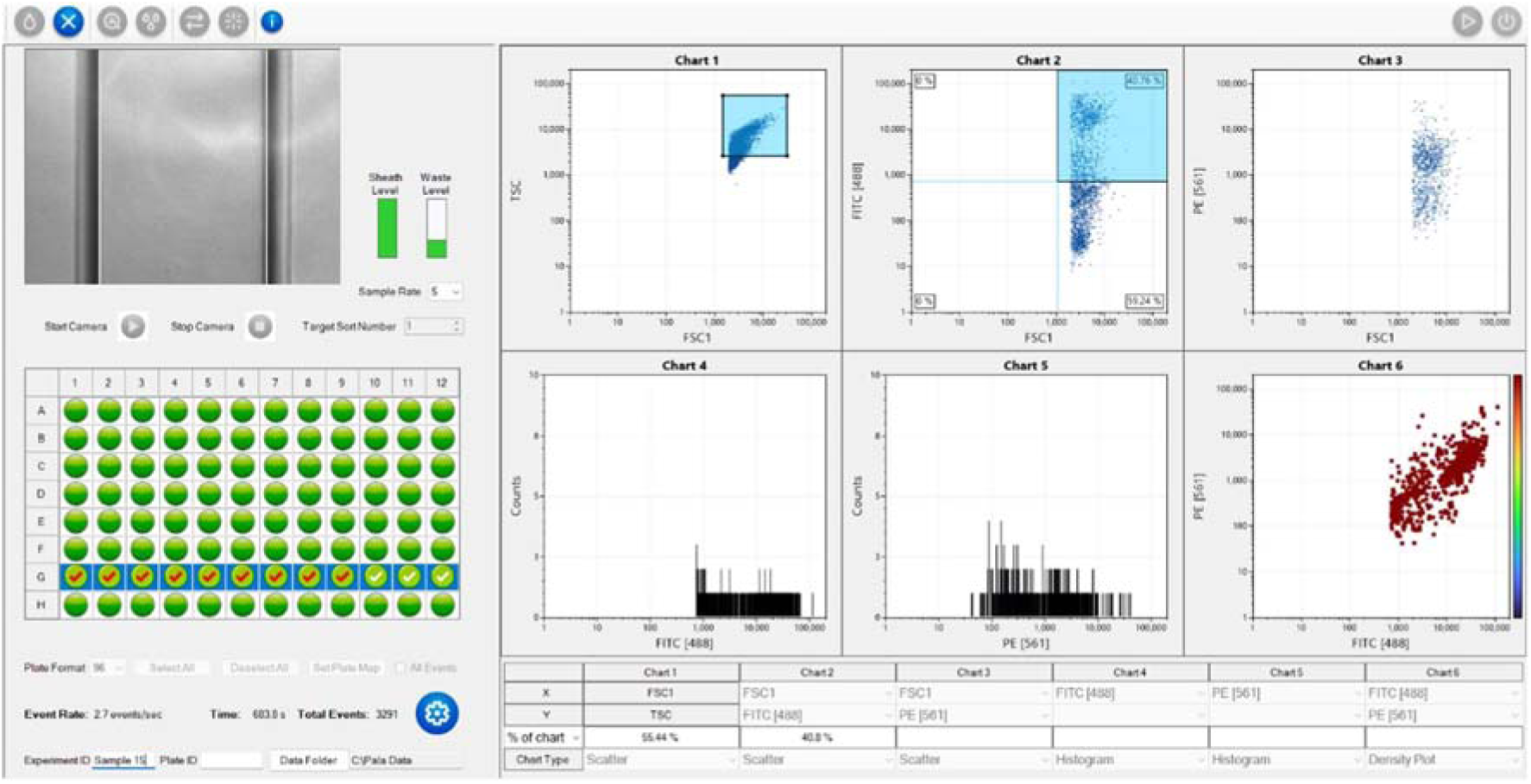
Single cell dispensing of Sample 5 (OCAS12). Sample density was ∼3 events/sec with the FSC trigger level set to 2000. FITC+ only were selected, with a large number of dispensed events observed as also being PE+. Fifty target cells were dispensed into 9 wells.

### b. Tumour-Spiked into Blood Sample

**Figure S18:**
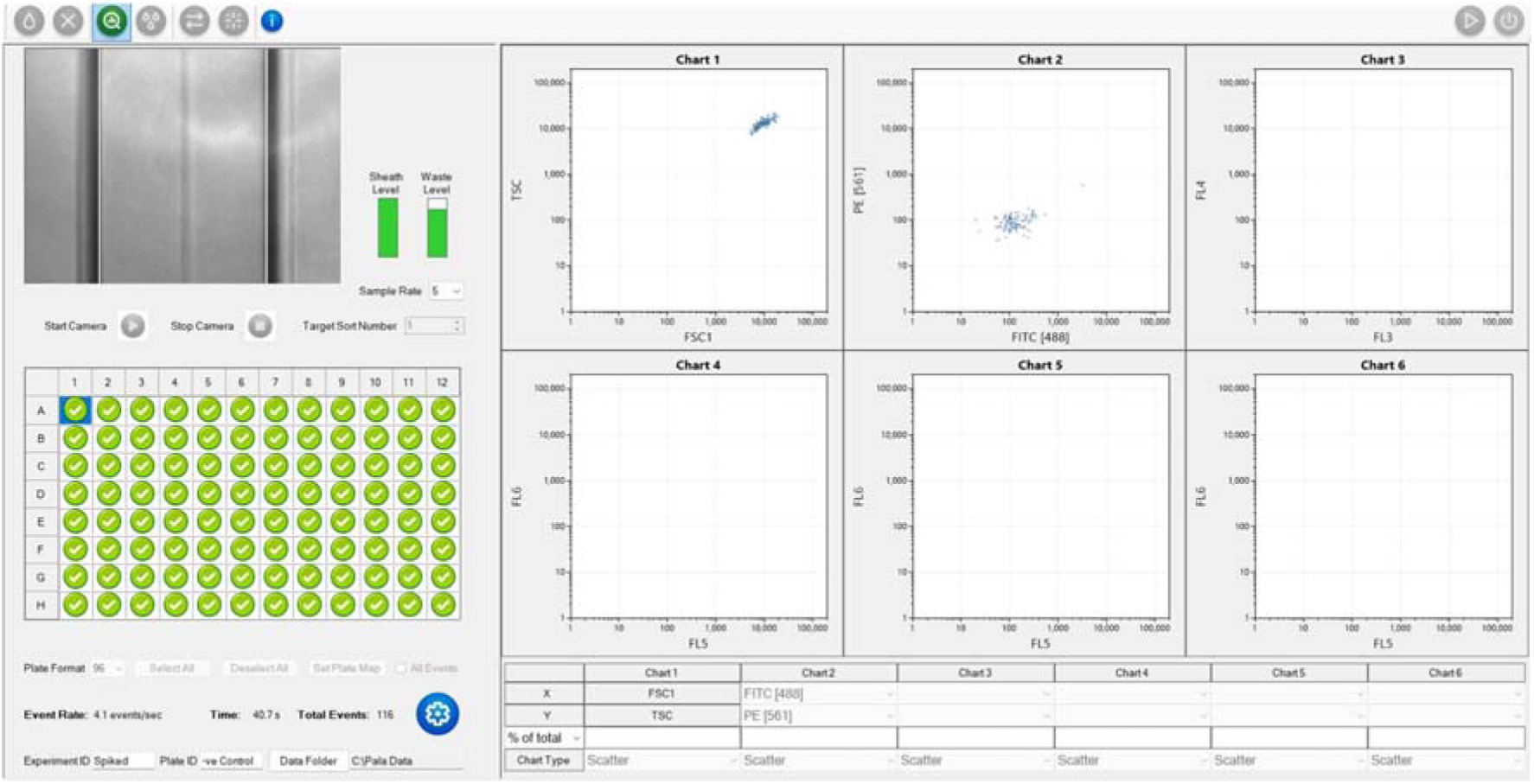
Unstained HeLa cell line to act as negative control sample. Sample density was ∼4 events/sec with the FSC trigger set to 5000.

**Figure S19:**
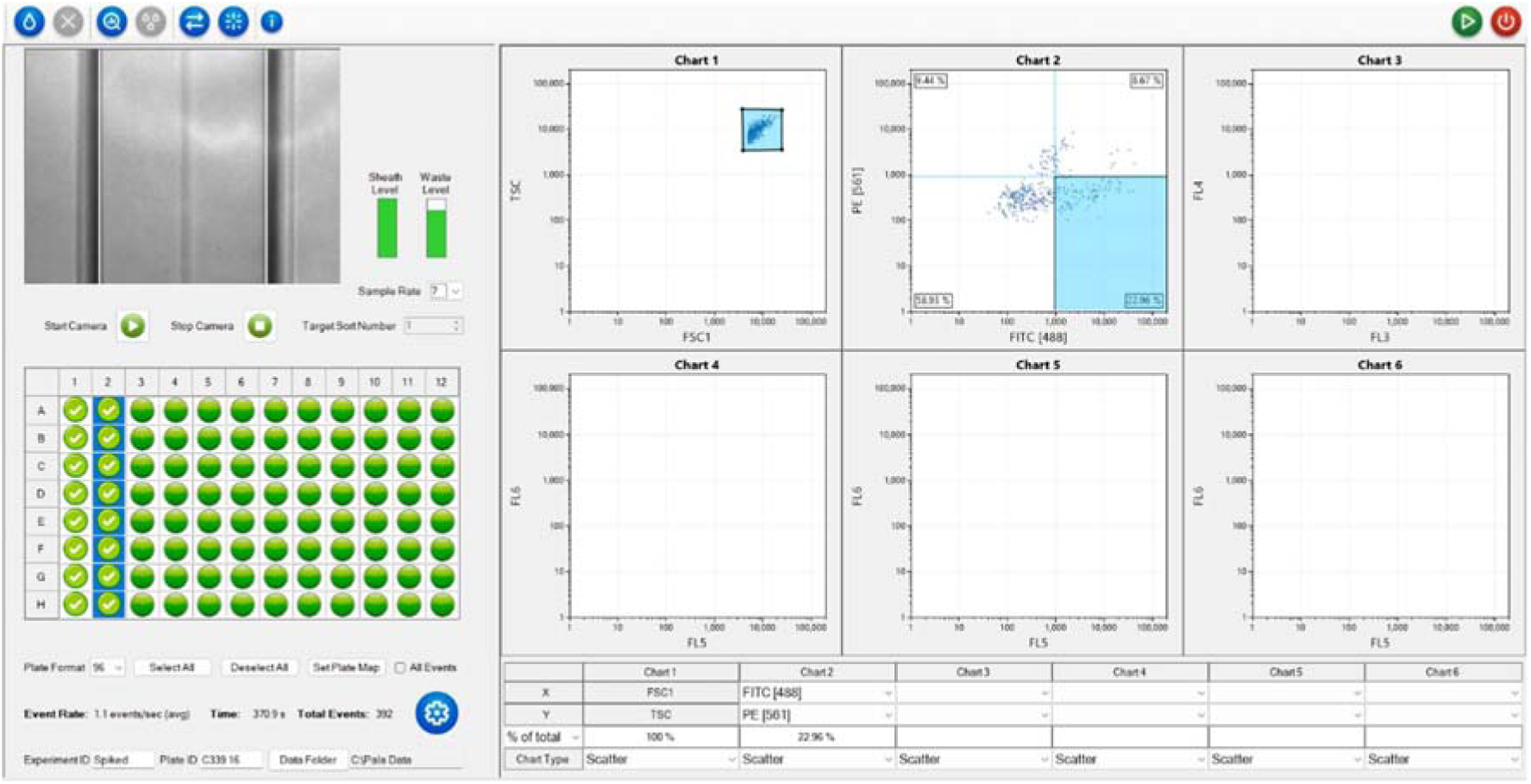
Single cell dispensing of spiked blood Sample 1 (C33a cells). Sample density was ∼1 event/sec with the fluorescent gate set to FITC 1000, and the FSC trigger set to 5000.

**Figure S20:**
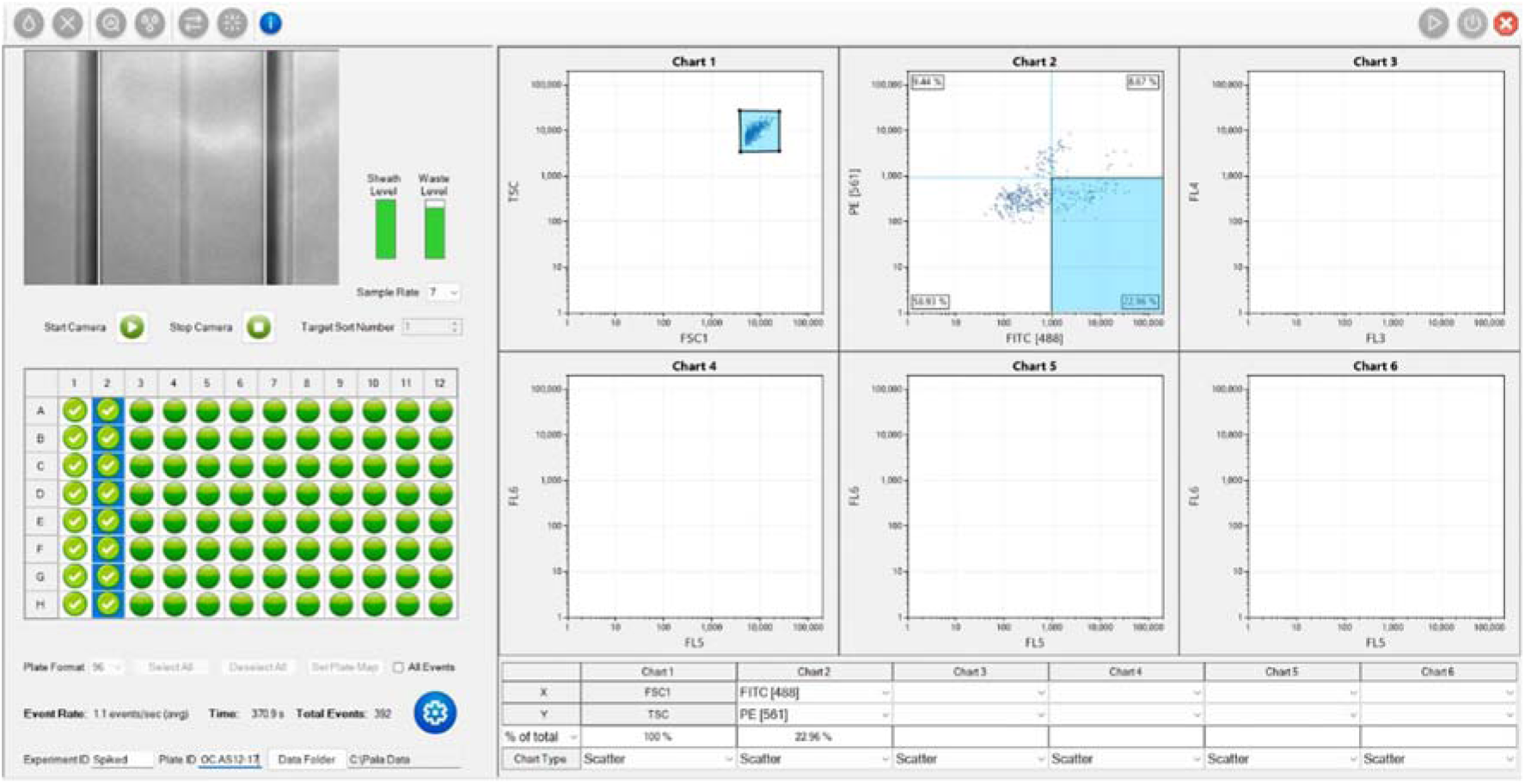
Single cell dispensing of spiked blood Sample 2 (OCAS12). Sample density was ∼1 event/sec with the fluorescent gate set to FITC 1000, and the FSC trigger set to 5000.

**Figure S21:**
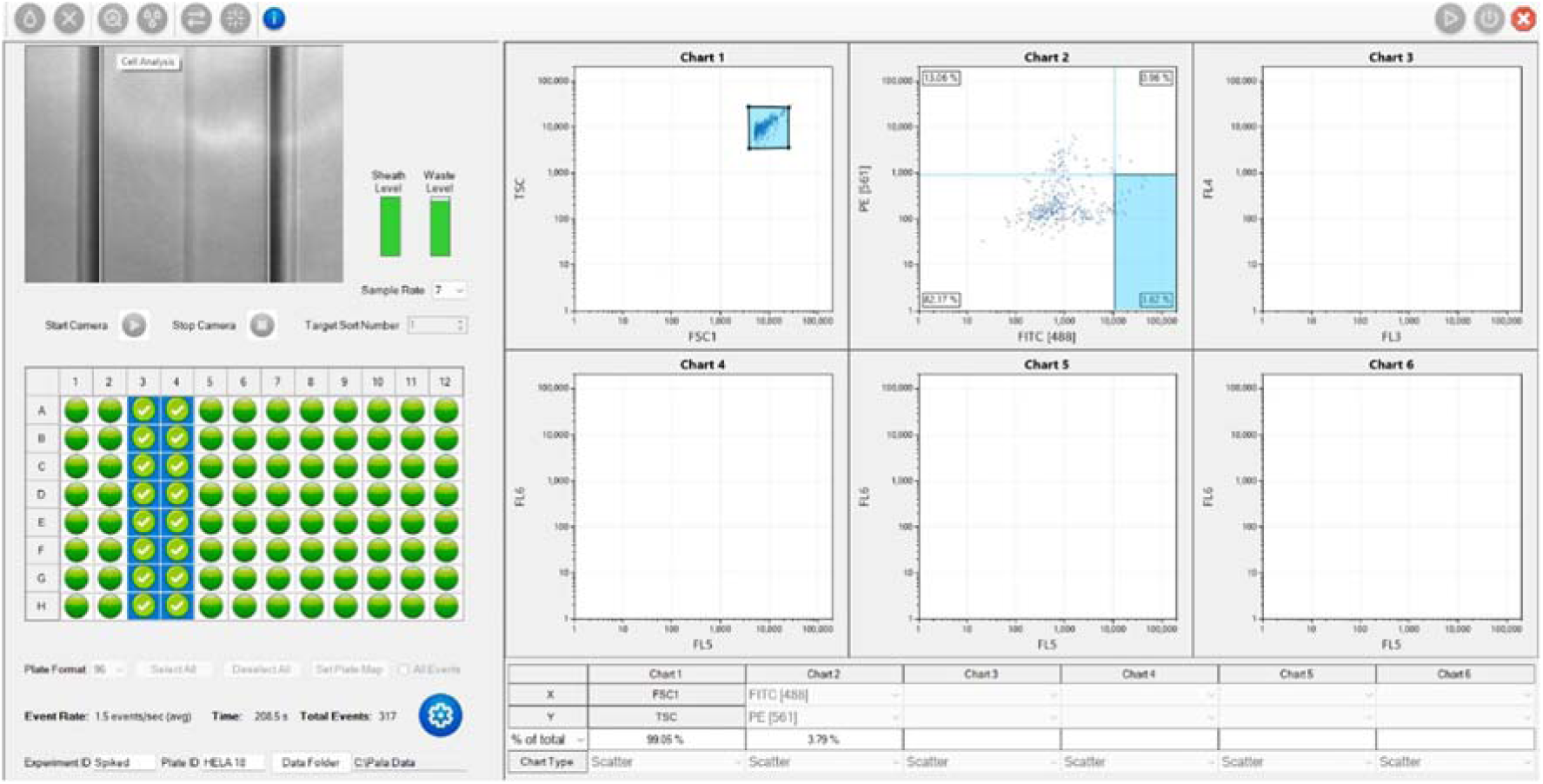
Single cell dispensing of spiked blood Sample 3 (HeLa cells). Sample density was ∼1 event/sec with the fluorescent gate set to FITC 10,000, and the FSC trigger set to 5000.

**Figure S22:**
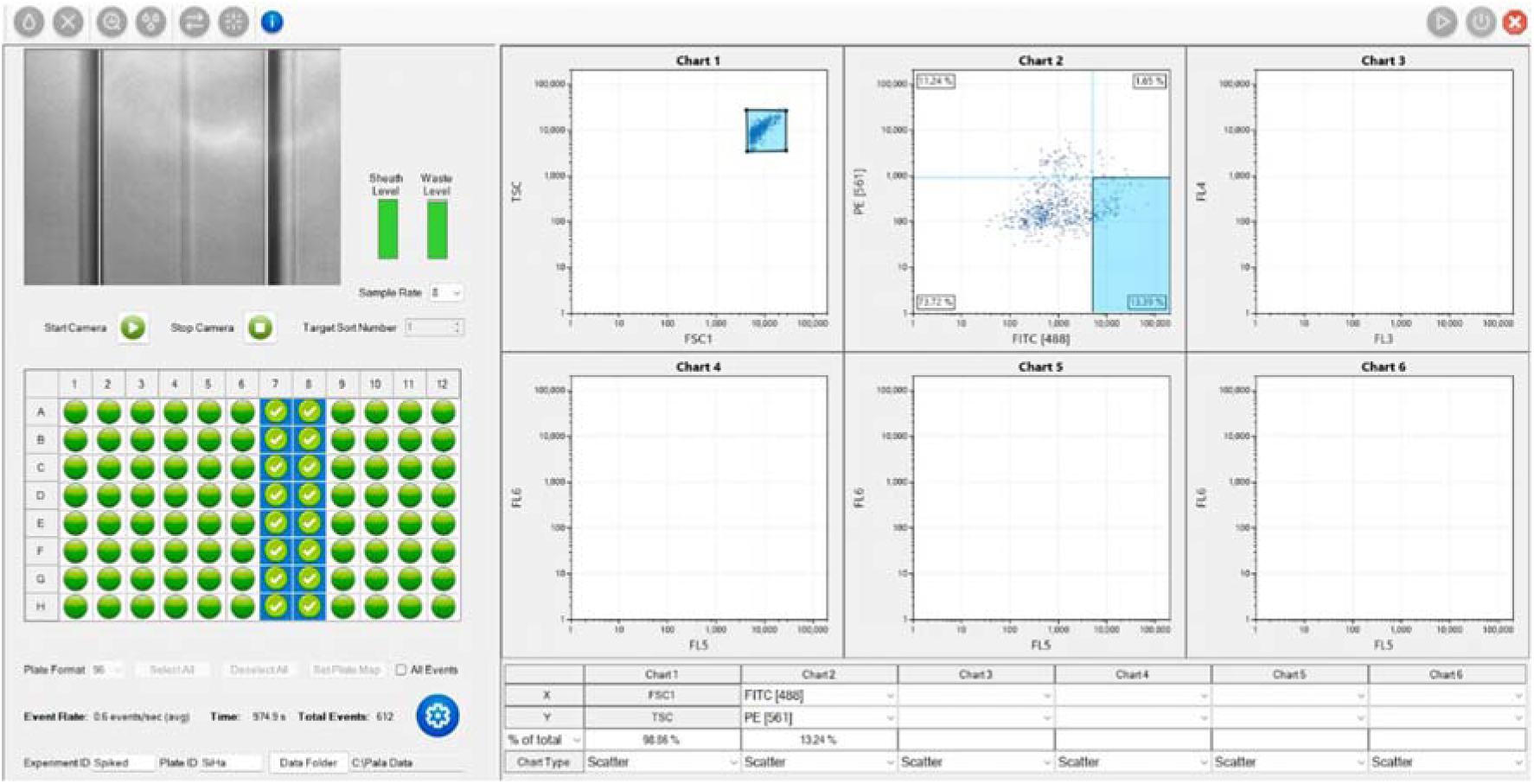
Single cell dispensing of spiked blood Sample 4 (SiHa). Sample density was ∼1 event/sec with the fluorescent gate set to FITC 5000, and the FSC trigger set to 5000.

**Figure S23:**
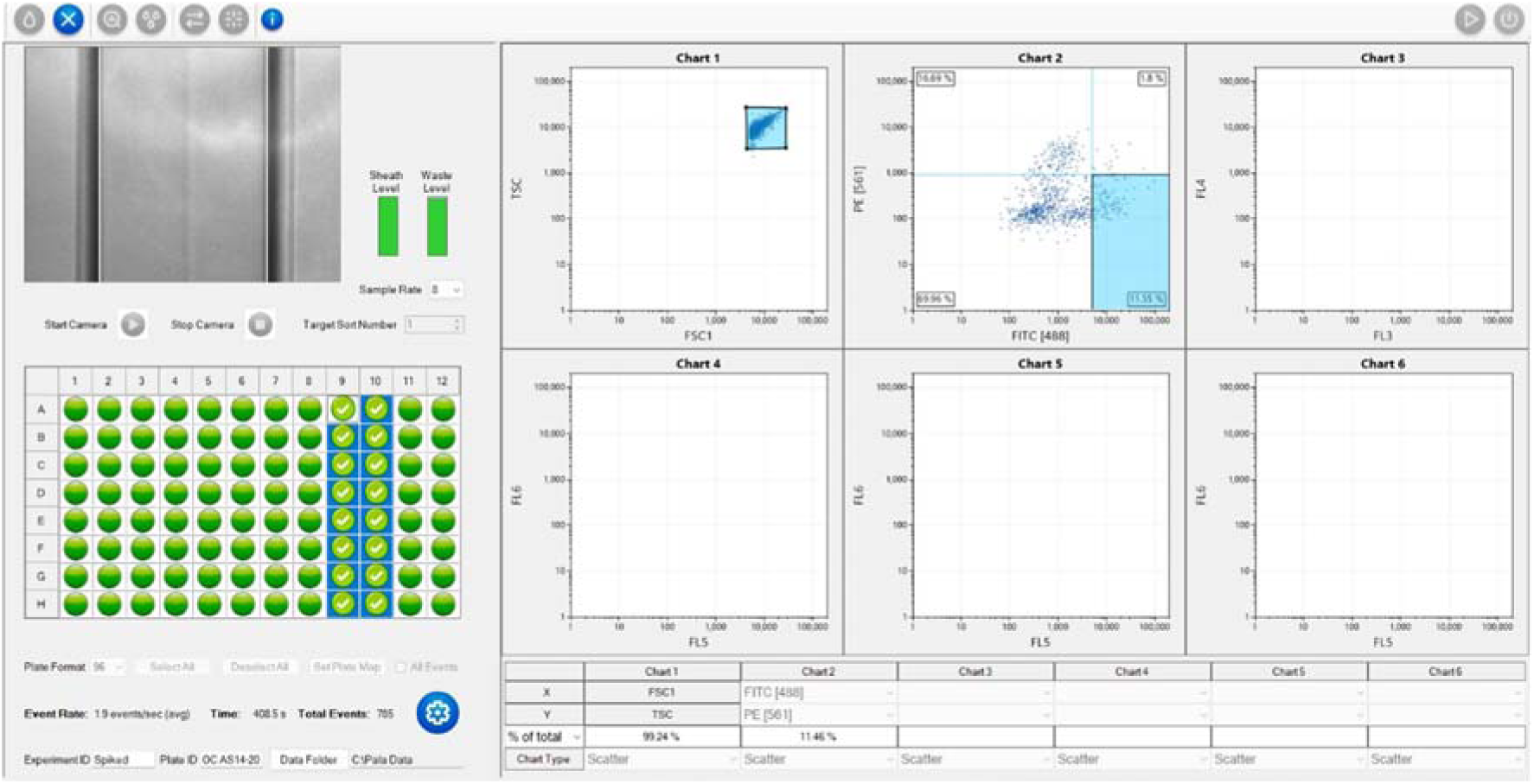
Single cell dispensing of spiked blood Sample 5 (OCAS14). Sample density was ∼2 events/sec with the fluorescent gate set to FITC 5000, and the FSC trigger set to 5000.

## 4. Patient-Derived Organoid Generation

**Figure S24:**
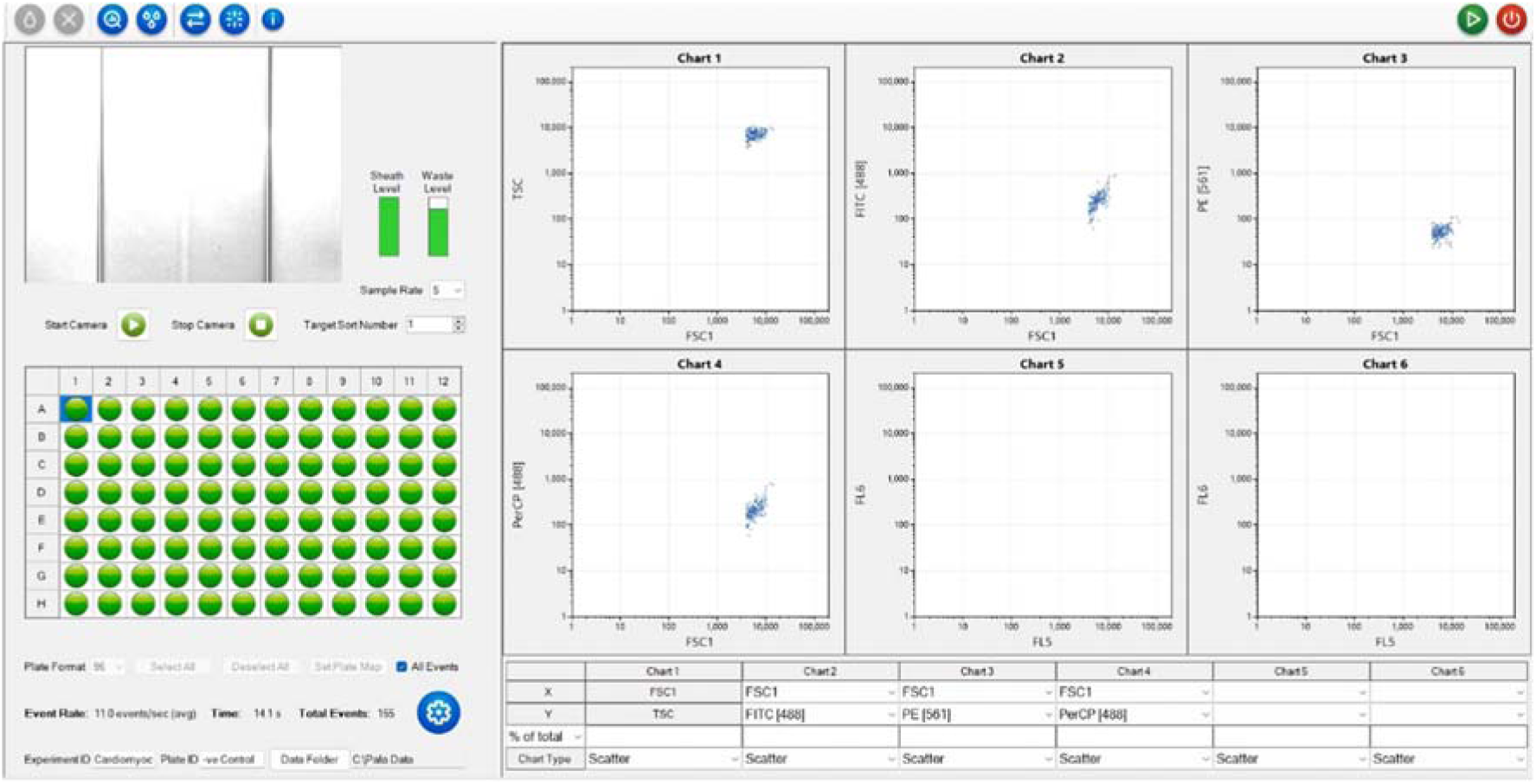
Single cell dispensing of organoid Sample 1 (OCAST16). Sample density was ∼2 events/sec with the FSC trigger level set to 7000.

**Figure S25:**
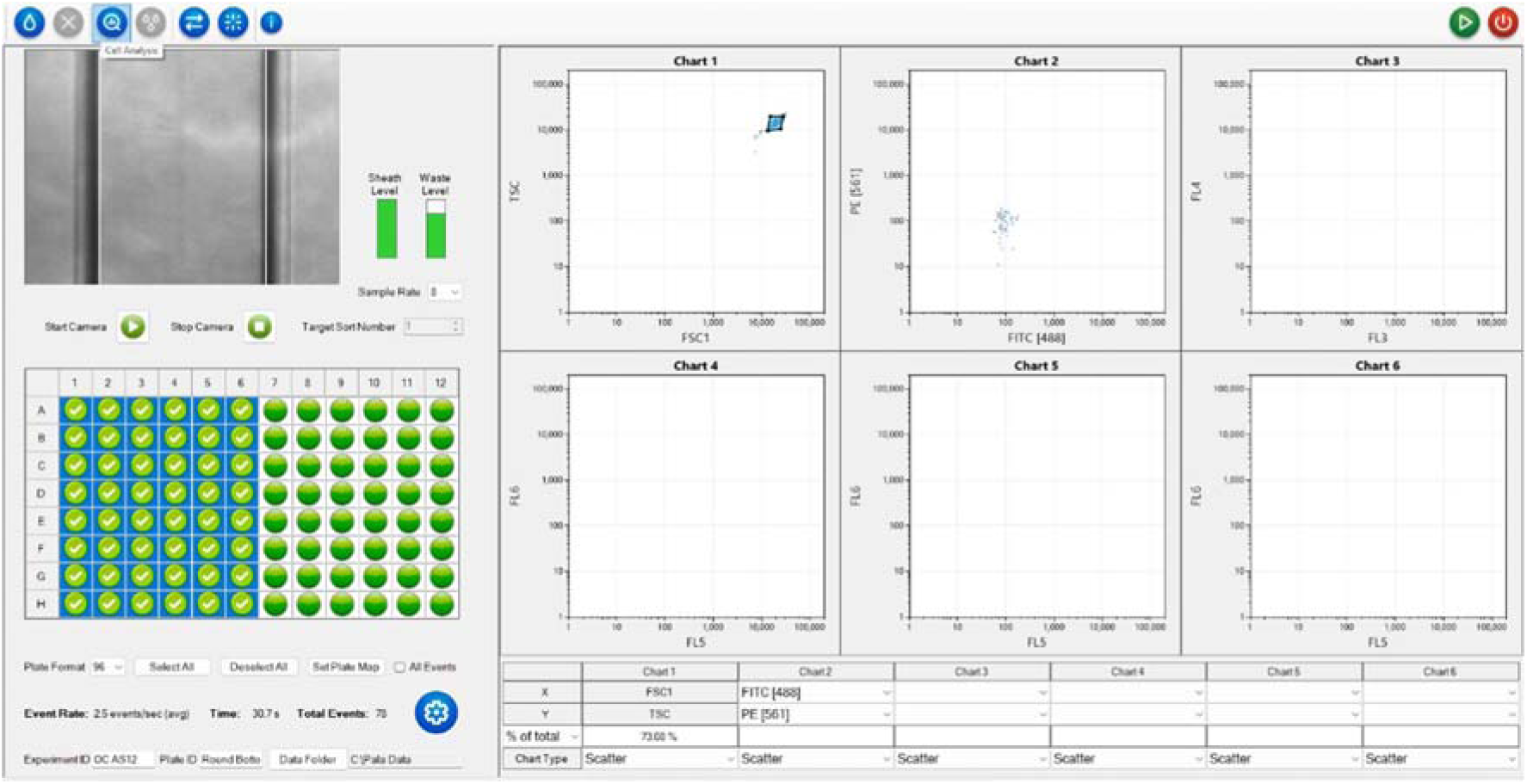
Single cell dispensing of organoid Sample 2 (OC AS12). Sample density was ∼2.5 events/sec with the FSC trigger level set to 7000.

**Figure S26:**
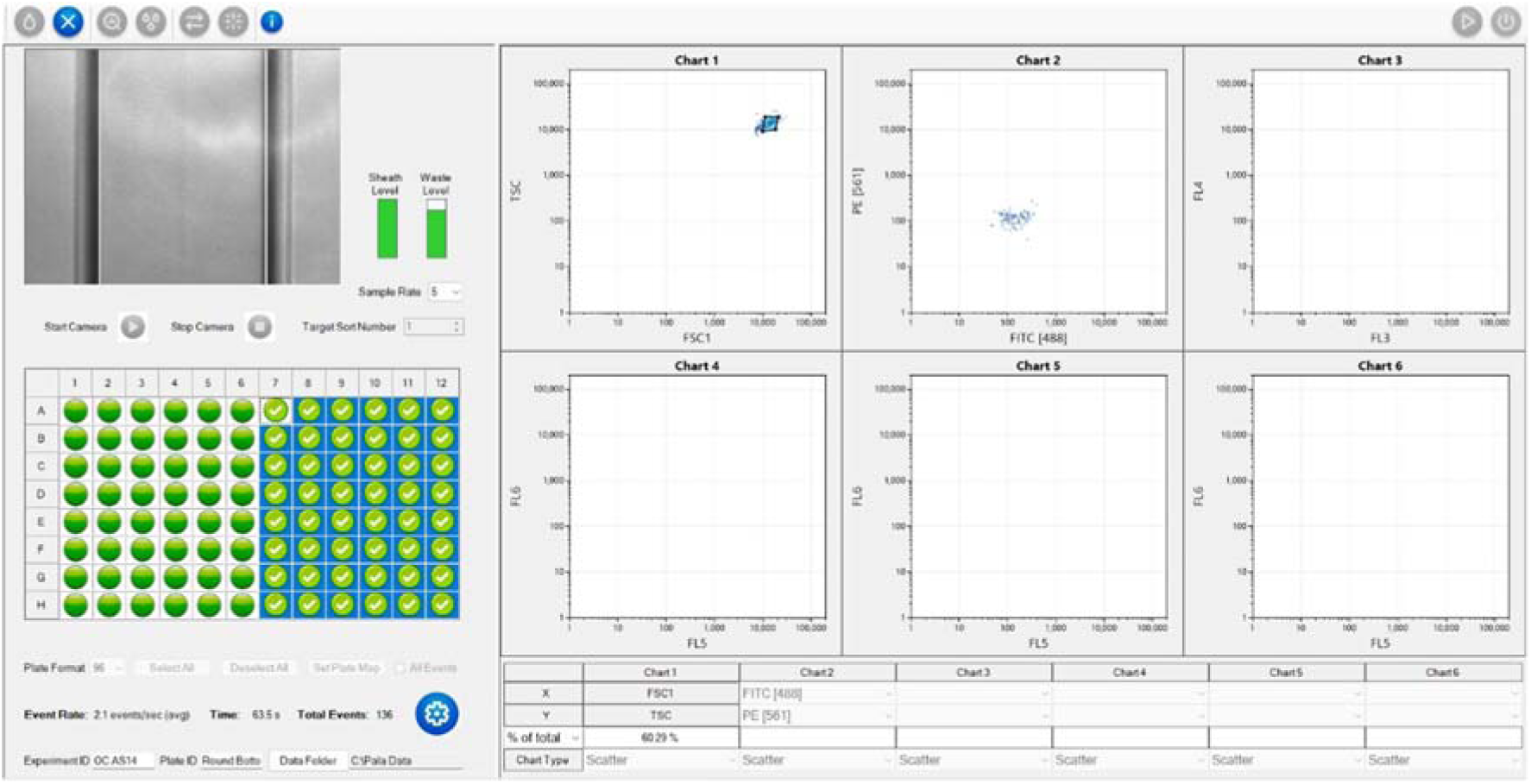
Single cell dispensing of organoid Sample 3 (OCAS14). Sample density was ∼2.5 events/sec with the FSC trigger level set to 7000.

